# Noncanonical activation of protease-activated receptor 1 induces autophagy in mammalian breast cancer cells

**DOI:** 10.1101/2025.01.11.632575

**Authors:** Saibal Saha, Rishikesh Kumar, Snehanjan Sarangi, Animesh Gope, Ananda Pal, Rima Tapader, Niraj Nag, Amit Pal

**Author notes:** Corresponding author Address: Division of Molecular Pathophysiology, ICMR-National Institute for Research in Bacterial Infections P-33, CIT Road, Scheme XM, Beliaghata, Kolkata-700010 West Bengal, India.

## Abstract

Whether autophagy is a boon or a bane for the body does not have a straightforward answer, as its consequences are intricately tied to the pathophysiological context that triggers its activation. The intricate nature of the effects of autophagy on the body, compounded by its multifaceted regulation through various upstream signaling pathways, warrants a relatively highly deep and nuanced research. This study delves into the regulation of autophagy through noncanonical activation of protease-activated receptor 1 (PAR1) by hemagglutinin protease (HAP). Unlike the canonical activation of PAR1 by thrombin, which enhances cell proliferation via mechanistic target of rapamycin (mTOR) signaling, HAP-induced activation downregulates mTOR and triggers autophagy in mammalian breast cancer cells. This noncanonical activation generates an N-terminal sequence in PAR1, which, when mimicked by a synthetic peptide, induces autophagy independently of HAP. Further investigation in BALB/c mouse model of low-grade breast cancer reveals that the synthetic peptide-induced autophagy significantly inhibits tumor growth and delays carcinogenesis progression. Importantly, the lack of PAR1 expression in normal, healthy cells facilitates the peptide to selectively target cancerous cells with relatively high PAR1 expression, highlighting its potential as a therapeutic tool against low-grade breast cancer. These findings provide valuable insights into autophagy modulation via PAR1 and suggest a promising avenue for targeted cancer therapy.

## Introduction

The literal meaning of “autophagy” as derived from its Greek origin (Greek word “Auto” means self and “Phagos” means eat) [1] does not completely encapsulate its multifaceted roles across various cellular contexts. Beyond its cytoprotective functions, macroautophagy (hereafter referred as autophagy) can lead to a specific form of cellular death known as type II programmed cell death. Autophagy is an evolutionarily conserved adaptive response of cells, initiated by a number of external and internal stimuli [2, 3]. Generally, cells recruit autophagy as a sophisticated recycling mechanism, in which dysfunctional or superfluous organelles, invading pathogens, or nonessential organelles under stressed condition are enclosed within a double-membrane vesicle termed phagophore, resulting in the formation of autophagosome (AP). Sequestosome 1 (SQSTM1/p62) and next to BRCA1 gene 1 protein act as adapter molecules for transporting selected ubiquitinated molecules to the phagophore [4–6]. AP subsequently fuses with lysosome to form autolysosome (AL) that degrades sequestered components through proteolytic activity of lysosomal hydrolases, and the building blocks are released to the cytosol for further utilization [7]. In incomplete autophagy, various stimuli initiate the process of phagophore formation and sequestration but fail to further proceed to the fusion of AP and lysosome, resulting into accumulation of cargo [8]. Impaired autophagy is implicated in several neurodegenerative disorders including Alzheimer’s and Parkinson’s diseases, cancers, autoimmune diseases, and metabolic syndromes including type 2 diabetes [9].

In cancer, autophagy plays a complex and dual role that varies with the disease stage. In the early stages of cancer, autophagy acts as a tumor-suppressive mechanism by degrading misfolded proteins and pathogens, potentially preventing further tumor development.

Conversely, in later stages, cancer cells exploit autophagy for surviving and adapting to the hostile environment within the host [10–13].

The interplay between proteasomal degradation and autophagy serves as a compensatory mechanism for the breakdown of various cellular components; an obstruction in one pathway often augments the other. For instance, inhibition of 26S proteasome leads to the activation of AMP-activated protein kinase (AMPK) [14–16], a positive regulator of autophagy [17]. Nevertheless, certain proteases facilitate the induction of autophagy. HsApg4A, a cysteine protease, cleaves the C-terminal end of the three mammalian homologs of autophagy-related protein (Atg) 8 (Gamma-aminobutyric acid receptor-associated protein (GABARAP), Microtubule-associated protein light chain 3 (LC3), and Golgi-associated ATPase enhancer of 16 kDa (GATE-16)), thereby promoting the formation of autophagic vacuoles [18, 19].

Proteases can trigger various signaling pathways by interacting with protease-activated receptors (PARs), a class of G protein-coupled receptors, which include four subtypes (PAR1–4) [20]. Among them, the expression of PAR1, PAR2, and PAR4 is significantly higher in cancer cells than in normal cells, making them promising targets for cancer therapy [21]. Thrombin activates PAR1 by making a proteolytic cleavage at its N-terminal extracellular region, and the N-terminal region acts as a tethered ligand, which binds to the second extracellular loop of the receptor. This thrombin-mediated activation of PAR1 upregulates the phosphorylation of mechanistic target of rapamycin (mTOR) [22], a master regulator of autophagy [23, 24]. Therefore, the canonical activation of PAR1 by thrombin activates coagulation cascades and leads to several other cellular events, including induction of angiogenesis and promotion of growth and proliferation of metastatic cancer cells [25].

Hemagglutinin protease (HAP), a 35-kDa zinc-dependent secretory metalloprotease, isolated from *Vibrio cholerae* O1 strain (C6709), induces apoptosis in mammalian breast cancer cells [26]. Among the four subtypes of PARs, HAP specifically engages with PAR1, facilitating its activation through proteolytic cleavage at the N-terminus. In Ehrlich ascites carcinoma (EAC) cells, a murine breast cancer cell line, this cleavage generates a new N-terminal extracellular segment, sequenced as PFISEDASGY. Albeit, the precise region of this newly formed N-terminal end, which functions as a tethered ligand and intramolecularly binds to the receptor for initiating downstream signaling pathways, remains unidentified [27]. Subsequently, based on the sequence of the HAP-mediated cleavage site of PAR1 in EAC cells, a peptide (PFISED) has been designed, which induced apoptosis in both breast and colon cancer cell lines [28].

Evidently, autophagy being an early response of cells and apoptosis being a late one [29], they share some common upstream signaling pathways, allowing a single stimulus to activate both the processes [30–32]. However, the regulation of autophagy through noncanonical PAR1 activation by HAP remains largely unexplored. Therefore, the current study explored the potential of HAP-mediated noncanonical activation of PAR1 to induce autophagy in a dose-and time-dependent manner in mammalian breast cancer cells.

## Materials and Methods

### Bacterial strain growth conditions

The *Vibrio cholerae* O1 strain C6709 was selected for this study due to its harboring of a functional hapR gene, which encodes a regulatory protein that modulates several cellular processes in *Vibrio cholerae*, including the expression of hemagglutinin HAP [33]. Routine cultivation of *Vibrio cholerae* (C6709) was carried out in Luria broth (Himedia, M575) at 37[with shaking for 18 h [34]. To preserve bacterial stocks, cells were stored at-70[in Luria broth supplemented with 20% sterile glycerol (Sigma-Aldrich, G5516) [33].

### Purification of the protease

HAP was isolated from *Vibrio cholerae* O1 strain C6709 and purified according to the protocol outlined by Ghosh *et al*. (2016) [35]. The *V. cholerae* O1 strain C6709 was grown and harvested as previously described. The cell-free culture supernatant was sequentially filtered through 0.45-μm and 0.22-μm cellulose acetate membranes (Merck, Darmstadt, Germany, Product No. HAWP04700 and Product No. GSWP04700) to eliminate residual cells. Outer membrane vesicles were subsequently removed from the filtrate by ultracentrifugation at 150,000 x g for 3 h at 4 using a T-865 rotor (Sorval Instruments, USA) [36]. The supernatant was carefully aspirated and subjected to salting out with 60% saturated ammonium sulfate (Sigma-Aldrich, St. Louis, Missouri, USA, Product No. A4418). Protein precipitation was achieved by centrifugation at 10,000 rotation per minute (RPM) for 20 min at 4. The resulting pellet was resuspended in 25 mM Tris (hydroxymethyl) aminomethane hydrochloride buffer (pH 7.4) (Sigma-Aldrich, St. Louis, Missouri, USA, Product No. 648317), dialyzed against the same buffer, and concentrated using an Amicon® Ultra Centrifugal Filter (Millipore, UFC9050). The protein was then purified through a single-step ion-exchange chromatography column (2.5 × 20 cm) packed with Diethylaminoethyl cellulose-52 (DEAE-52) matrix (Whatman, 4057050) and pre-equilibrated with 25 mM Tris (hydroxymethyl) aminomethane hydrochloride buffer (pH 7.4). Proteins were eluted in the unbound fractions, which formed distinct peaks, pooled, dialyzed, concentrated, and assayed for protease activity. Notably, the purification of HAP was performed without the use of Ethylenediamine tetraacetic acid (EDTA). Chromatographic separation was conducted using a Biologic Duo Flow Chromatographic System (Bio-Rad, USA). The purified HAP was stored in aliquots at-20.

### Sodium Dodecyl Sulfate-Polyacrylamide Gel Electrophoresis (SDS-PAGE)

SDS-PAGE was performed using the method described by Laemmli [37]. Protein samples were denatured in sample buffer and separated on a 12.5% polyacrylamide gel using a discontinuous buffer system.

### Protease activity assay

In this study, azocasein (Sigma-Aldrich, A2765) was employed as a substrate to assess the proteolytic activity of HAP [38]. The substrate-enzyme mixture (azocasein-HAP) was incubated at 37[for 1 hour, followed by termination of the reaction by the addition of 10% (w/v) trichloroacetic acid (Sigma-Aldrich, T8657). The precipitated protein was eliminated by centrifugation at 12,000 RPM for 4 min, and the supernatant was transferred to a clean tube containing 525 mM sodium hydroxide. Absorbance was measured at 440 nm using a Shimadzu spectrophotometer (Shimadzu, Japan). Negative controls included the substrate with buffer and the substrate with EDTA-inactivated enzyme.

### Cell culture

Human epithelial breast cancer cells MCF7 (ATCC, HTB-22), murine epithelial breast cancer cells 4T1 (ATCC, CRL-2539), and human normal breast epithelial cells MCF10A (ATCC, CRL-10317) were cultured (passages 10-12) and maintained in Minimum Essential Medium (MEM) (Gibco, 61100061), RPMI 1640 (Gibco, 23400021) and Mammary Epithelial Cell Basal Medium (MEBM) (Lonza, CC-3151) Mammary Epithelial Cell Growth Medium (MEGM) SingleQuots supplements (Lonza, CC-4136) respectively. Each medium was supplemented with 10% Fetal bovine serum (FBS) (Gibco, 10270106), 100 U/ml penicillin G, and 100 µg/ml streptomycin sulfate (Gibco, 15140122) and cells were incubated at 37[in a 5% CO2 atmosphere using a Heracell incubator (Thermo Fisher Scientific, Waltham, MA, Catalog Number 51026331). The MCF7 and MCF10A cell lines were generously provided by Dr. Somsubhra Nath, Presidency University, Kolkata, while the 4T1 cell line was a gift from Dr. Saptak Banerjee, CNCI, Kolkata.

### Western blotting

Western blotting was conducted as previously described by Nag *et al*. [39]. In brief, cultured cells were lysed using Radioimmunoprecipitation assay (RIPA) buffer (Sigma-Aldrich, R0278), and protein concentrations were determined using the Bicinchonic acid (BCA) assay kit (Thermo Fisher Scientific, 23225). Equal amounts of protein were loaded onto SDS-PAGE for separation and subsequently transferred electrophoretically to a Polyvinylidene fluoride (PVDF) membrane (Merck, IPVH00010). Following transfer, the membrane was blocked with 3% Bovine serum albumin (BSA) (Sigma-Aldrich, A1470) in Tris-buffered saline (TBS) and incubated overnight at 4[with the primary antibody (diluted according to the manufacturer’s instructions) against the target protein. The blot was then washed with Tris-buffered saline-Tween 20 (TBST) buffer and incubated with Horseradish peroxidase (HRP)-conjugated secondary antibody (Cell Signaling Technology, 7076; 1:1000 and Cell Signaling Technology, 7074; 1:1000) for 2 h at room temperature. Protein bands were visualized using a Bio-Rad gel documentation system with an Enhanced chemiluminescence (ECL) substrate (Thermo Fisher Scientific, 34580) specific for HRP-conjugated antibodies. Western blot quantification was performed using GelQuant (Thermo Fisher Scientific, USA) and ImageJ (NIH, Bethesda, MD) software. At least three independent experiments were conducted to validate the results, and band intensities were normalized to loading controls: GAPDH for MCF7 and MCF10A cells, and Cyclophilin B for 4T1 cells.

### Analyses of lysosomal and autophagosomal vacuoles by confocal microscopy

Lysosomal and autophagosomal vacuoles were analyzed by staining with LysoTracker™ Green DND-26 (Thermo Fisher Scientific, L7526), following the method described by Klepka *et al*. [40] with minor modifications tailored to the cell lines. Briefly, MCF7 and 4T1 cells were seeded onto cover slips in 6-well plates at a density of 0.3 × 10^5^ cells per well.

After 24 h of incubation, cells were treated with HAP (0.5 μg/ml for MCF7 and 0.75 μg/ml for 4T1) for 30 min, with or without pre-treatment with 25 mM 3-MA, an autophagy inhibitor (Sigma-Aldrich, M9281) for 24 h. Untreated cells and those treated only with 3-MA served as normal and negative controls, respectively. Following HAP treatment, cells were fixed with 4% paraformaldehyde for 10 min at room temperature. They were then permeabilized with 0.1% Triton X-100 (Sigma-Aldrich, T8787) in 0.1% sodium citrate solution and blocked with 5% FBS prepared in TBST. The cells were incubated with LysoTracker™ Green DND-26 for 45 min at 37[. Nuclei were stained with Hoechst 33342(Cell Signaling Technology, 4082; working concentration 1.0 μg/ml) for 10 min at 37[. After washing twice, cells were mounted with 10% glycerol (Sigma-Aldrich, G5516). Images were acquired using a confocal microscope (Zeiss, Germany, LSM 710) with a Plan-Apochromat 63X / 1.40 NA objective. Data were processed and analyzed using Zen 3.4 (blue edition) and ImageJ software.

### Autophagy detection by confocal microscopy and flow cytometry

Autophagy was detected by confocal microscopy using the Autophagy Assay Kit (Abcam, ab139484) according to the manufacturer’s instructions. The kit contains a green fluorescent dye that specifically stains autophagosomal and autolysosomal vacuoles. Briefly, MCF7 and 4T1 cells were seeded onto cover slips in 6-well plates at a density of 0.3 × 10^5 cells per well. After 24 h of incubation, cells were treated with HAP (0.5 μg/ml for MCF7 and 0.75 μg/ml for 4T1) for 30 min and with peptides (50μM dosage of PAFISED for MCF7 and 75 μM dosage of PFISED for 4T1) for 4 h. Following treatment, cells were washed twice with 1X Assay Buffer (diluted from the 10X stock provided with the kit) supplemented with 5% FBS. Cells were then incubated at 37[for 30 min with 100 μl of staining solution, which contained 2 μl of the green fluorescent dye and 1 μl of nuclear stain (Hoechst), both provided in the kit, in 1 ml of 1X Assay Buffer, supplemented with 5% FBS. After incubation, the staining solution was removed, and cells were washed twice with 1X Assay Buffer containing 5% FBS. Cells were then fixed with 4% formaldehyde for 15 min and washed thrice with 1X Assay Buffer containing 5% FBS. Finally, cells were mounted on glass slides with 10% glycerol. Green fluorescence was detected using the FITC filter, and nuclear staining was visualized using the DAPI filter on a confocal microscope (Zeiss, Germany, LSM 710) with a Plan-Apochromat 63X / 1.40 NA objective. Data were processed and analyzed using Zen 3.4 (blue edition) and ImageJ software.

The induction of autophagy was further analyzed by flow cytometry using the same Autophagy Assay Kit following the manufacturer’s protocol. In a brief, cells were treated with HAP and the peptides as mentioned earlier and harvested by brief incubation with 0.25% trypsin-EDTA (Gibco, 25200072) at 37[. Cells were washed with 1X Assay Buffer, supplemented with 5% FBS, as outlined earlier and stained with 250 µl of the staining solution, comprising of 1 μl of the green fluorescent dye (provided in the kit) in 1 ml of 1X Assay Buffer supplemented with 5% FBS. Flowcytometry data were acquired using BD FACSAria™ II (BD, USA). Analyses of the flow cytometry data was performed using FloJo™ 10 software.

### Immunocytochemistry and confocal imaging

Immunocytochemistry and confocal imaging were conducted following the procedures outlined by Nag *et al*. [39]. In brief, MCF7 and 4T1 cells were grown on coverslips in 6 well cell culture plates and treated with HAP (0.5 µg/ml for MCF7 and 0.75 µg/ml for 4T1) for 30 min. Following the treatment, cells were fixed with 4% paraformaldehyde for 10 min at room temperature. Cells were subsequently permeabilized using 0.1% Triton X-100 in 0.1% sodium citrate solution and blocked with TBST supplemented with 5% FBS for 2 h. They were incubated overnight at 4[in a moist chamber with primary antibody (diluted according to the manufacturer’s instructions) against the protein of interest. Following primary antibody incubation, cells were washed with TBS and incubated for 2 h at room temperature with Alexa 488 conjugated (Cell Signaling Technology, 4408; 1:200 and Cell Signaling Technology, 4412; 1:500) or Alexa 555 conjugated (Cell Signaling Technology, 4413; 1:500 and Cell Signaling Technology, 4409; 1:500) secondary antibodies. Nuclei were stained with Hoechst 33342 (Cell Signaling Technology, 4082; working concentration 1.0 μg/ml) for 10 min at 37[. After washing the cells twice with TBS, they were mounted with 10% glycerol.

Confocal images were captured using a confocal microscope (Zeiss, Germany, LSM 710) with a Plan-Apochromat 63X / 1.40 NA objective. Image processing and analyses were performed using Zen 3.4 (blue edition) and ImageJ software.

### Raising of antisera against purified HAP

Antiserum against HAP was generated by immunizing a New Zealand White rabbit, following the method described by Mondal *et al*. [41]. The rabbit was initially administered an intramuscular injection of 100 mg of purified HAP emulsified with an equal volume of Freund’s complete adjuvant (Sigma-Aldrich, F5881). This was followed by four booster injections of 100 mg of HAP emulsified with Freund’s incomplete adjuvant (Sigma-Aldrich, St. Louis, Missouri, USA, Product No. F5506) at 7-day intervals. Blood samples were collected from the rabbit on day 0 and three days after the final booster injection, allowing the blood to clot overnight at room temperature. The sera were then collected, centrifuged (1,000 RPM for 5 min), diluted in autoclaved glycerol, and stored at-80 until use at a 1:1000 dilution.

### Colocalization analyses by confocal imaging

Colocalization of HAP with PAR1 and of LAMP1 with LC3II was analyzed in HAP-treated (0.5 µg/ml for MCF7 and 0.75 µg/ml for 4T1 for 30 min) and peptide-treated (50 µM of PAFISED for MCF7 and 75 µM of PFISED for 4T1 for 4 h) cells, respectively, by confocal microscopy, following the protocol described by Das *et al*. [42]. In brief, cells were grown on coverslips in 6 well cell culture plates. After the treatment, they were fixed with 4% paraformaldehyde for 10 min at room temperature, and then permeabilized with 0.1% Triton X-100 in 0.1% sodium citrate solution. The cells were blocked with 5% FBS in TBST for 2 h and incubated overnight at 4[in a moist chamber with both of primary antibodies (diluted as per the manufacturer’s instructions) against the respective proteins of interest. After washing twice with TBS, cells were incubated for 2 h at room temperature with Alexa 488 (Cell Signaling Technology, 4408; 1:500 and Cell Signaling Technology, 4412; 1:500) and Alexa 555 (Cell Signaling Technology, 4413; 1:500 and Cell Signaling Technology, 4409; 1:500) conjugated secondary antibodies. Nuclei were stained with Hoechst 33342 (Cell Signaling Technology, 4082; working concentration 1.0 μg/ml) for 10 min at 37[. Following two washes with TBS, cells were mounted with 10% glycerol on a glass slide. Confocal images were acquired using a Zeiss LSM 710 microscope (Zeiss, Germany) with a Plan-Apochromat 63X / 1.40 NA objective. Image processing and analyses were performed using Zen 3.4 (blue edition) and ImageJ software. Colocalization was quantified by calculating Pearson’s correlation coefficient.

### Knock down of PAR1

MCF7 cells were first starved for 18 h in incomplete MEM before transfection. To transiently knock down PAR1, PAR1 siRNA (CGGUCUGUUAUGUGUCUAUdTdT) (Eurogentec, Seraing, Belgium) was used. Lipofectamine 3000 (Thermo Fisher Scientific, L3000001) served as the transfection reagent, and the procedure was carried out according to the manufacturer’s instructions. Transfection was performed when the cells reached approximately 70% confluency. The transfection reagents were diluted in Opti-MEM (Gibco, 31985062) and added onto the cells dropwise. After 4 h, the medium was replaced with fresh complete MEM (supplemented with 10% FBS). PAR1 siRNA was transfected for 72 h to achieve efficient knockdown. Scrambled siRNA (Santa Cruz Biotechnology, sc-37007) was used as a control in this knockdown study. Knockdown efficiency was confirmed by Western blotting.

### Flow cytometric analyses of lysosomal and autophagosomal vacuoles in PAR1 knocked down cells

Flow cytometry was employed to analyze lysosomal and autophagosomal vacuoles in HAP-treated WT and PAR1 KD MCF7 cells. Detection of these vacuoles was achieved by staining with LysoTracker™ Green DND-26 (Thermo Fisher Scientific, L7526), following the protocol outlined by Chikte *et al*. [43] with cell-specific optimization of dye concentration and incubation time. WT and PAR1 KD MCF7 cells were seeded at a density of 0.3 × 10^5^ cells per well in 6-well tissue culture plates and were grown overnight. Both WT and PAR1 KD cells were treated with HAP at dosages of 0.5 μg/ml and 0.75 μg/ml for 30 as well as 60 min. After treatment, cells were harvested by trypsinization (0.25% Trypsin-EDTA), washed twice with TBS, and resuspended in 500 μl of TBS. Cells were incubated with 60 nM LysoTracker™ Green DND-26 for 45 min at 37[for staining followed by washing with TBS twice. Finally, cells were resuspended in 500 µl of TBS and Flow cytometric analyses were conducted on the single cell populations, gated by forward scatter (FSC) and side scatter (SSC). The FITC channel (excitation at 488 nm and detection range of 515-545 nm) was used to identify LysoTracker™ Green DND-26-positive cells. Data acquisition was performed using a BD FACSAria™ II (BD, USA) and analyses were carried out using FloJo™ 10 software.

### in silico analysis

The crystal structure of human PAR1 bound to Vorapaxar (PDB ID: 3VW7) was retrieved from Protein Data Bank (https://www.rcsb.org) [44]. The physio-chemical properties, drug-likeness and pharmacokinetics of peptide were checked using swissADME (https://www.swissadme.ch) and toxicity was checked by ToxinPred (https://crdd.osdd.net/raghava/toxinpred) [45, 46]. After checking the peptides properties, three servers were used one for peptide structure modelling and other two were used for molecular docking. Servers were PEP-FOLD4 (https://bioserv.rpbs.univ-paris-diderot.fr/services/PEP-FOLD4), Autodock Vina (https://www.swissdock.ch), CABS-dock (https://biocomp.chem.uw.edu.pl/CABSdock) [47–50]. PEP-FOLD4 was used for the peptide structure prediction; Autodock Vina and CABS-dock were used for the molecular docking in which Blind docking was performed by CABS-dock. The docking study was conducted in the following steps: - 1. Re-docked Vorapaxar by Autodock Vina [51], 2. Peptide docking against PAR 1 by Autodock Vina 3. Blind docking by CABS-dock 4. Binding interactions and analysis was performed by Biovia Discovery Studio visualizer 2024 (Dassault Systèmes BIOVIA, USA).

### Transmission electron microscopy (TEM) imaging

For the TEM experiment, HAP and peptide treated and untreated MCF7 and 4T1 cells were washed three times in phosphate-buffered saline (PBS) and fixed with 2.5% glutaraldehyde in PBS (pH 7.4) overnight at 4[. The samples were then dyed with 2% uranyl acetate for 30[min, washed three times with distilled water and dehydrated for 15 min. Subsequently, the samples were embedded in EPON 812 resin (Head Biotechnology, E8000) and cured for 24[h at 70[. Ultrathin slices were procured using an ultramicrotome and then subjected to staining with a solution containing 2% uranyl acetate and lead citrate. The samples were imaged with a FEI Tecnai 12 BioTwin Transmission Electron Microscope operating at 100[kV.

### Analysis of autophagy flux by pMXs GFP-LC3-RFP reporter construct

Autophagic flux in MCF7 and 4T1 cells treated with HAP and peptides was assessed by transiently expressing the pMXs GFP-LC3-RFP plasmid in these cells. This plasmid was originally cloned in NEB® Stable Competent E. coli (New England Biolabs, USA) and obtained as an agarose stab culture as a generous gift from Noboru Mizushima Laboratory (Addgene, 117413) [52]. The plasmid was isolated using the QIAprep Spin Miniprep Kit (Qiagen, 27104) according to the manufacturer’s protocol, and its purity was confirmed by agarose gel electrophoresis.

Prior to transfection, MCF7 and 4T1 cells were cultured on coverslips and subjected to starvation for 18 h in their respective media. Transfection was performed using Lipofectamine 3000 (Thermo Fisher Scientific, L3000001), following the manufacturer’s protocol. Cells were transfected at approximately 70% confluency, and the all the reagents, including P3000, was added in Opti-MEM (Gibco, 31985062). After 4 h, the media was replaced with fresh complete media (supplemented with 10% FBS) suitable for the respective cell lines. MCF7 and 4T1 cells were transfected for 72 and 48 h, respectively, to ensure optimal transfection, which was confirmed by fluorescence microscopy. pAM_1C Empty Vector (Active Motiff, 53023) transfected cells were used as vehicle controls.

Following transfection, cells were treated with HAP (0.5 µg/ml for MCF7 and 0.75 µg/ml for 4T1) for 30 min or with peptides (50 µM dosage of PAFISED for MCF7 and 75 µM dosage of PFISED for 4T1) for 4 h, and then fixed with 4% paraformaldehyde. After fixation, cells were stained with Hoechst 33342 (Cell Signaling Technology, 4082; working concentration 1.0 μg/ml) for 10 min at 37 and mounted with 10% glycerol on glass slides. Images were acquired using a confocal microscope (Zeiss, Germany, LSM 710) with a Plan-Apochromat 63X / 1.40 NA objective. Data processing was carried out using Zen 3.4 (blue edition) software, and the number of red and green puncta was quantified and analyzed using ImageJ software.

### Cell viability and cell proliferation assay

MTT assay was conducted according to the standard protocol described by Kar *et al*. [53] to assess the effect of HAP and peptides (PAFISED for MCF7 and PFISED for 4T1 cells) on the cellular viability of breast cancer cells. Approximately 1 × 10^4^ viable cells per well were seeded in flat bottom 96-well cell culture plates, where viability was determined using 0.4% Trypan blue solution (Thermo Fisher Scientific, 15250061) before seeding. After 24 h of incubation, MCF7 and 4T1 cells were treated with the optimized concentrations of HAP (0.5 μg/ml for MCF7 and 0.75 μg/ml for 4T1) for 30 min and peptides (50 μM dosage of PAFISED for MCF7 and 75 μM dosage of PFISED for 4T1) for 4 h, with or without pre-treatment with 25 mM 3-MA, an autophagy inhibitor for 24 h. Following the treatment with HAP and the peptides, MTT solution (Thermo Fisher Scientific, M6494; 0.8 mg/ml in serum-free medium) was added, and cells were incubated for 4 h in the dark at 37. The media was then removed, and 100 µl Dimethyl sulfoxide (Sigma-Aldrich, D5879) was added in each well to solubilize the purple crystal formazan. After a 15-minute incubation in the dark, absorbance was measured at 595 nm using a spectrophotomteric plate reader. The mean of three replicates was used to calculate the cell survival percentage.

To examine the impact of HAP-and peptide-induced autophagy on breast cancer cell proliferation, Ki-67 nuclear antigen immunofluorescence assay was performed in MCF7 and 4T1 cells following the standard protocol [54] with slight modifications optimized for the current experimental condition. Cells were treated with HAP and the respective peptides (PAFISED for MCF7 and PFISED for 4T1) as described above. After treatment, cells were fixed with 4% paraformaldehyde for 10 min at room temperature, permeabilized with 0.1% Triton X-100 in 0.1% sodium citrate solution, and blocked with 5% serum in TBST. Cells were incubated overnight at 4[with Ki-67 (D3B5) Rabbit mAb (Cell Signaling Technology, 9129; 1:400) in a moist chamber. Following washing with TBS, cells were incubated with Alexa 488-conjugated anti-rabbit IgG antibody (Cell Signaling Technology, 4412; 1:200). Nuclei were stained with Hoechst 33342 (Cell Signaling Technology, 4082; working concentration 1.0 μg/ml) for 10 min at 37[. After two additional washes with TBS, cells were mounted on glass slides with 10% glycerol. Images were captured using a confocal microscope (Zeiss, Germany, LSM 710) with a Plan-Apochromat 63X / 1.40 NA objective. Data were processed and analyzed using Zen 3.4 (blue edition) and ImageJ software.

### *in vivo* experiments Animal model

In order to establish a low-grade breast tumor mouse model for studying the effect of the peptide (PFISED) induced autophagy in early stage breast cancer, first solid breast tumors were developed by implanting 4T1 cells in 4-6 weeks old female BALB/c mice following the protocol described by Pulaski *et al*. [55]. Briefly, Solid tumors were developed by implanting 4T1 cells (0.5×10^7^ cells/kg of body weight) subcutaneously at the abdominal mammary fat pad and allowed them to multiply. Cells were suspended in serum free RPMI 1640 media and each mouse was injected with 100 µl of the suspension. The mice were randomly distributed in four groups, each containing 5 mice (n=5). The groups were stratified based on the time which the tumors were harvested at after the implantation of 4T1 cells. (i) Group I: harvested the tumor at day 7, (ii) group II: harvested the tumor at day 14, (ii) group III: harvested the tumor at day 21, (iv) group IV: harvested the tumor at day 28.

Tumors from all the groups were scored on the basis of histopathological analyses and grades were determined following the Nottingham Grading System [56, 57]. They have been further correlated immunohistochemically and with the size of the tumors.

Before evaluating the effect of the peptide PFISED on low grade breast tumor in BALB/c mice, the optimum dose of the peptide was identified based on the toxicity evaluation of the peptide following the protocol described by Lamichhane *et al*. [58]. Briefly peptide PFISED of six different doses (1, 2.5, 5, 7.5, 10 and 20 mg/kg body weight) was administered at an interval of 3 days starting from Day 0 (1^st^ dose) to Day 12 (5^th^ dose) and at day 15, blood was collected from retro-orbital sinus behind the eye. For the evaluation of hematological parameters, blood was collected in EDTA coated tubes. Hematological parameters including eosinophil, hemoglobin, lymphocyte, monocyte and neutrophil were counted using Medonic M32 Cell Counter (Boule, Sweden, M32B). Liver function was assessed by measuring ALT and AST in blood serum by ALT Activity Assay Kit (Sigma-Aldrich, MAK052) and Aspartate Transaminase (AST) Assay Kit (Sigma-Aldrich, MAK467) respectively and kidney function was evaluated by measuring creatinine and BUN (calculated from serum urea) by Creatinine Assay Kit (Sigma-Aldrich, MAK475) and Urea Assay Kit (Sigma-Aldrich, MAK471) respectively.

Based on the gradation of malignancy, out of the three groups with low-grade tumor, group II (carrying tumor of 14 days) was selected randomly for further analyses to evaluate the role of the peptide (PFISED) induced autophagy in low-grade murine breast cancer. Three groups of experimental mice were designed, each group containing 5 mice, carrying breast tumor of 14 days. (i) Group I: without any treatment, considered as control, (ii) group II: injected with 5 mg/kg of body weight dosage (as optimized by toxicity assessment of the peptide) of the peptide PFISED, (ii) group III: injected with the aforementioned dosage of peptide PFISED along with autophagy inhibitor, 3-MA (15 µg/gram of body weight). Mice of group II were injected with the peptide PFISED subcutaneously at the site of tumor at an interval of 3 days starting from the 14^th^ day to 26^th^ day after the implantation of 4T1 cells. In the mice of group III, 3-MA along with the peptide PFISED was administered according to the same experimental design of group II. The peptide PFISED and 3-MA were diluted in serum free RPMI 1640 and at one time each mouse was injected with 100 µl of the solution. Since, *in silico* analyses indicated that the peptide PFISED doesn’t follow the Lipinski and Veber rules, the possibility of oral administration of the peptide had been excluded.

All the animal experiments and animal study design were approved by the institutional animal ethics committee (IAEC) of ICMR-National Institute of Research in Bacterial Infection (formerly ICMR-NICED) (License No: PRO/168/Jan 2022). Mice were procured from the animal breeding facility of ICMR-National Institute of Research in Bacterial Infection and were supplied with food pellets and autoclaved water ad libitum. When required, all the mice were euthanized via asphyxiation in a CO_2_ chamber following the Committee for the Purpose of Control and Supervision of Experiments on Animals (CPCSEA) guidelines and the tumors were surgically excised.

### Histology and immunohistochemistry

Immunohistochemical and hematoxylin-eosin (H&E) staining of tumor tissues was done as per protocol described by Kim *et al*. [59] with some minor modifications as required for the experiment. In a brief, tissues were fixed in 10% formaldehyde for at least 24 h and then processed by an automated tissue processor (Leica TP1020, Leica Microsystems, Germany). Processed tissues were embedded in paraffin and sectioned (5 µM thin) from the paraffin block by microtome, deparaffinized using xylene (Sigma-Aldrich, 6.08685), serially rehydrated and permeabilized with 0.5% Triton X-100 (Sigma-Aldrich, 93443) for 10 min. Tissues were then incubated with 3% Hydrogen peroxide (H_2_O_2_) to inactivate the endogenous peroxidase activity and boiled in 10mM sodium citrate buffer pH 6.0 for 10 min antigen retrieval. Tissue sections were blocked by incubating with TBST containing 5% FBS for 1 hour. They were stained with hematoxylin and eosin for histopathological analyses and for immunohistochemical analyses they were incubated with Ki-67 antibody (Cell Signaling Technology, 12202; 1:200) overnight in a humidified chamber at 4, followed by an incubation with Alexa 488 conjugated goat anti-rabbit IgG antibody (Cell Signaling Technology, 4412) for 2 h at room temperature. Nuclei were stained with Hoechst.

Hematoxylin and eosin (H&E)-stained tissue sections were viewed under 20X magnification of Zeiss Axiovert 40 C microscope (Zeiss, Germany). Immunohistochemical images of randomly selected fields were captured using confocal microscope (Zeiss, Germany, LSM 710); objective-Plan-Apochromat 40X / 1.40 NA and quantified using Image J software.

## Statistical analyses

All experiments were replicated at least three times (n≥3). All animal experimental groups contained 5 animals each. The experimental results were presented as mean ± standard error of mean (SEM). All statistical analyses were done by applying ordinary one-way ANOVA or two-way ANOVA or Student’s t-test (Mann-Whitney test) or one sample t-test using GraphPad Prism 10. In all the graph panels, ns (non-significant) p ≥0.05, * (significant) 0.01≤ p <0.05, ** (very significant) 0.001≤ p <0.01, *** (extremely significant) 0.0001≤ p <0.001, **** (extremely significant) p<0.0001.

### Use of artificial intelligence

ChatGPT-4 has been employed to modify and improve the quality of language in some parts of the manuscript followed by rigorous manual revision.

### List of key materials

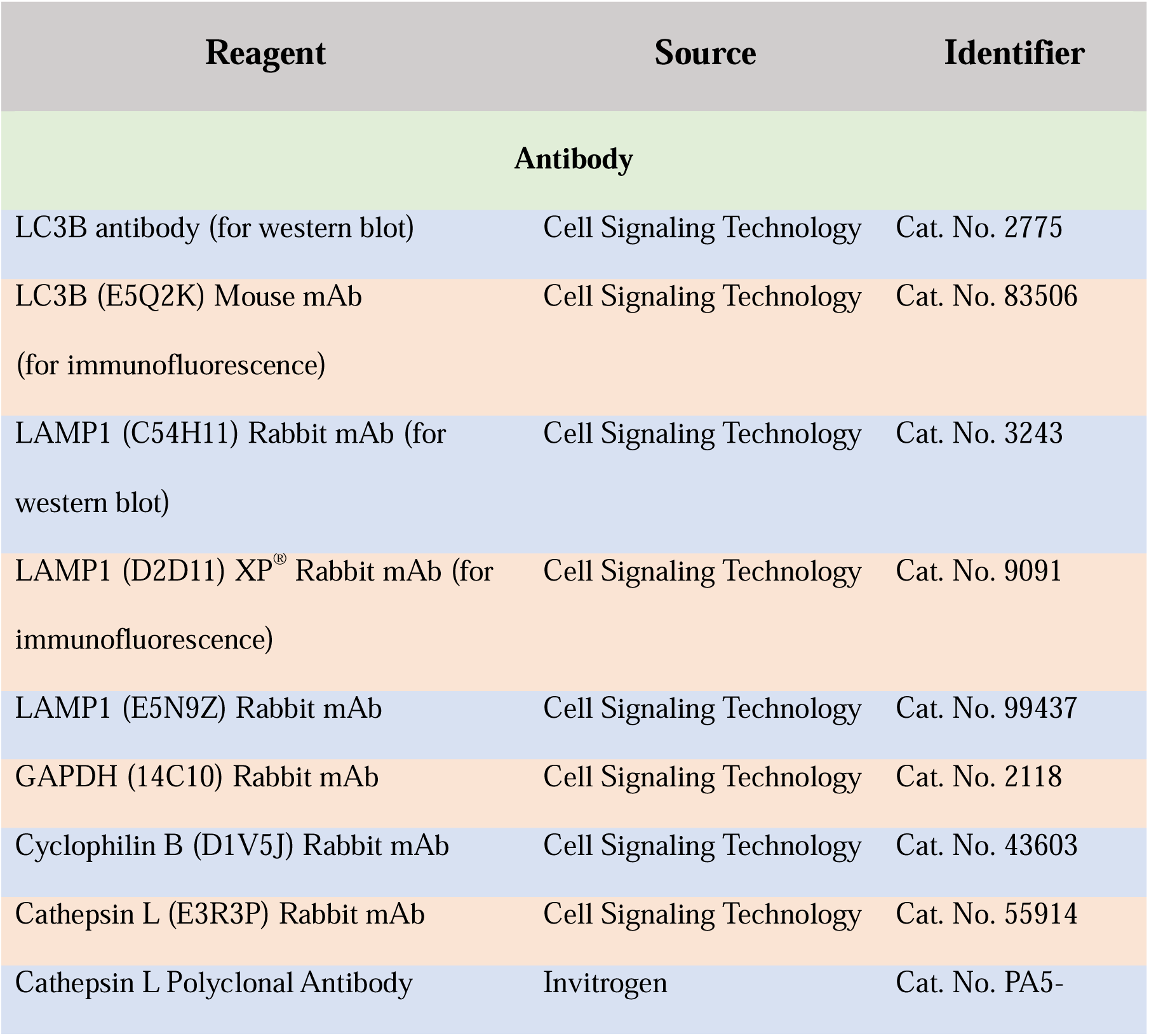
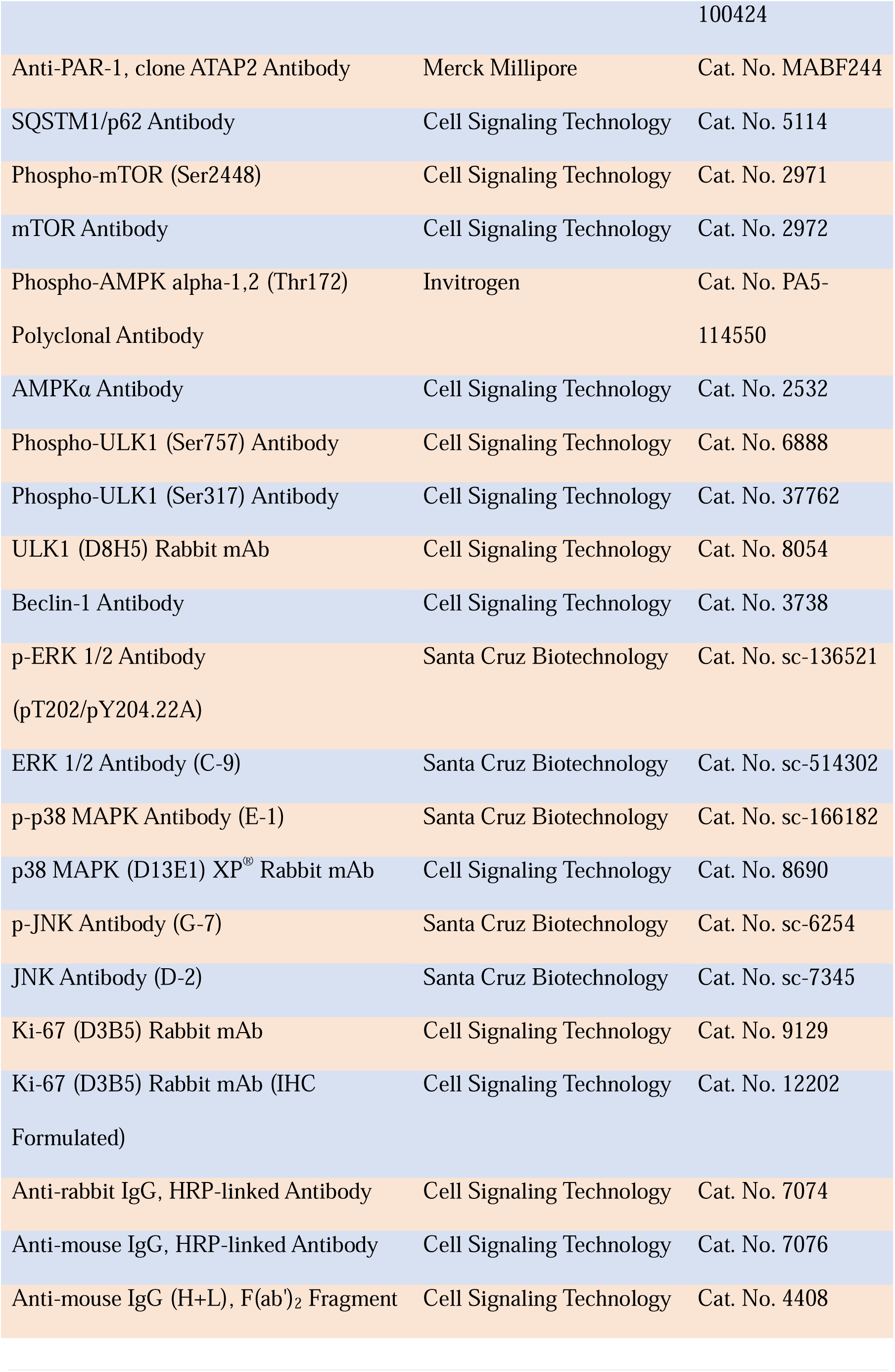
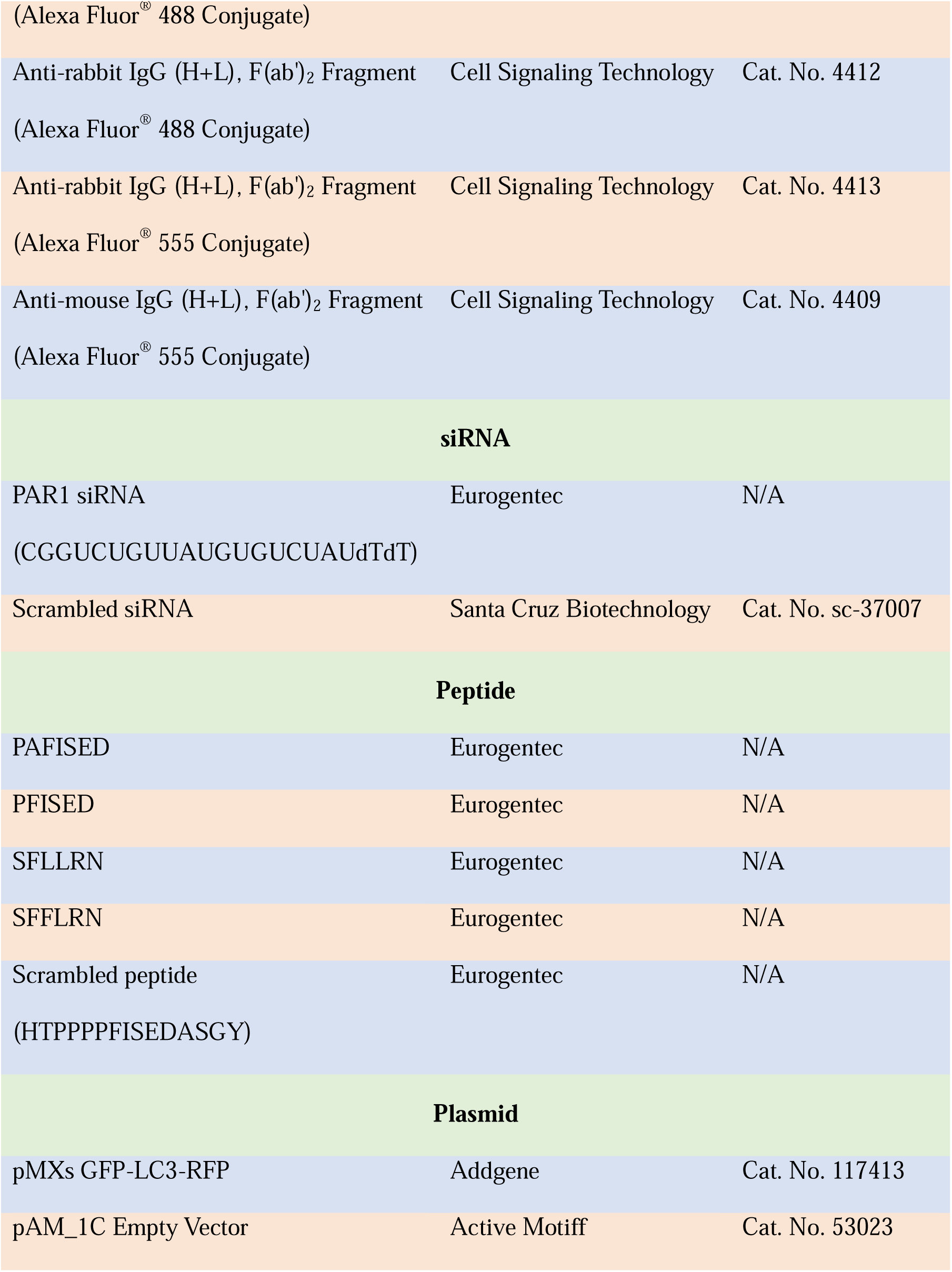

## Results

### HAP-induced autophagy

Immunoblot analysis was conducted to assess the expression of autophagy markers LC3II and Lysosome associated membrane protein (LAMP1) in human (MCF7, estrogen receptor+) and murine (4T1, estrogen receptor-) breast cancer cells treated with 0.25–1 µg/mL HAP for 30 or 60 min (Fig. 1Ai). Densitometric analysis revealed dose-and time-dependent increases in their expression. In MCF7 cells, optimal expression occurred with 0.5 µg/mL HAP treatment for 30 min, showing 3-and 22-fold increases for LC3II and LAMP1 expression, respectively. In 4T1 cells, LC3II and LAMP1 expression increased by 5-and 1.5-fold, respectively, with 0.75 µg/mL HAP treatment for 30 min. After 60 min, peak expression in 4T1 cells occurred with 0.5 µg/mL HAP, whereas MCF7 cells showed peak LAMP1 expression with 0.25 µg/mL HAP and LC3II expression with 0.5 µg/mL HAP (Fig. 1Aii). Based on these findings, HAP dosages were standardized to 0.5 µg/mL for MCF7 and 0.75 µg/mL for 4T1 for subsequent experiments.

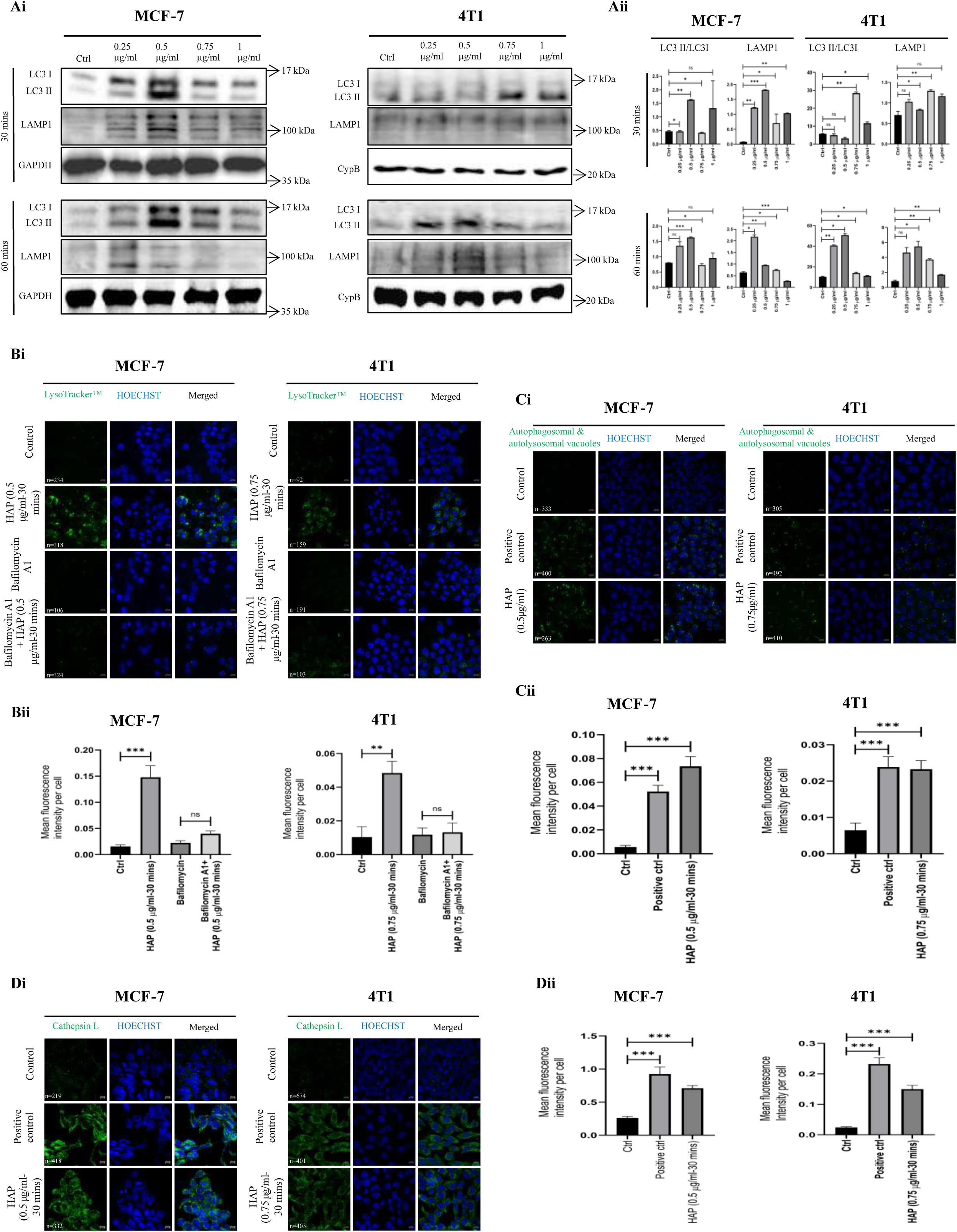

Confocal fluorescence microscopy showed increased lysosomal activity (Fig. 1Bi and Bii) and an elevation in the number of APs and ALs (Fig. 1Ci), as evident from the quantification of mean fluorescent intensity (MFI), indicating 13-and 3.6-fold increases in MCF7 and 4T1 cells, respectively (Fig. 1Cii). Pretreatment with bafilomycin A1 inhibited lysosomal accumulation, which was used as a negative control for lysosomal activity. Cells pretreated with rapamycin (500 nM) and chloroquine (60 μM) for 24 h were considered as positive controls for autophagy detection. Flow cytometric analysis of the HAP-treated cells for autophagy detection corroborated these findings (Supplementary Fig. 1Ai and Aii) revealing a significant increase in MFI (Supplementary Fig. 1Aiii).

Immunofluorescence studies for cathepsin L expression confirmed the ablation of autolysosomal membrane in HAP-treated MCF7 and 4T1 cells at the specified dosages and times (Fig. 1Di). Quantification of fluorescence revealed 2.7-and 6.1-fold higher expression of cathepsin L in the cytosol of HAP-treated MCF7 and 4T1 cells, respectively, than in their respective controls (Fig. 1Dii).

### PAR1 mediates HAP-induced autophagy

In MCF7 cells, PAR1 expression increased by 0.5 and 0.75 μg/mL HAP treatment for 30 min, and by 0.25 and 0.5 μg/mL HAP treatment for 60 min. In 4T1 cells, the highest upregulation occurred by 0.75 μg/mL HAP treatment for 30 min, and by 0.25 and 0.5 μg/mL HAP treatment for 60 min (Fig. 2Ai and Aii). These dosages aligned with those optimized for autophagy induction, suggesting a potential role of PAR1 in HAP-induced autophagy.

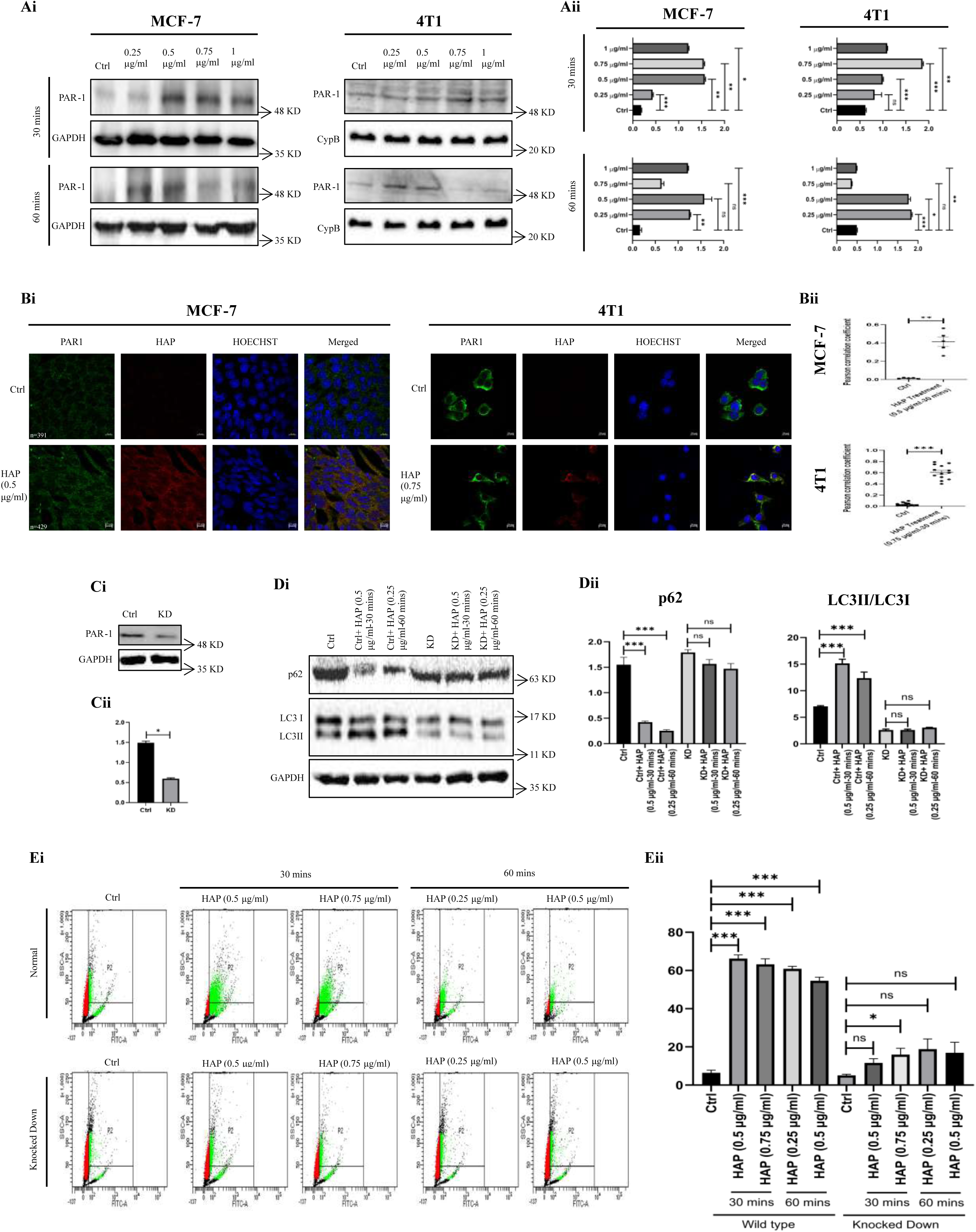

To investigate the interaction between HAP and PAR1, we used confocal microscopy to assess co-localization in cells treated with optimized HAP dosages for 30 min (Fig. 2Bi). Significant co-localization of HAP and PAR1 was noticed, with Pearson’s correlation coefficients indicating approximately 30-and 18-fold increases in MCF7 and 4T1 cells respectively, compared to those of controls (Fig. 2Bii).

To assess the role of PAR1 in autophagy, we transiently knocked down *PAR1* in MCF7 cells (Fig. 2Ci and Cii). HAP treatment (0.5 and 0.25 µg/mL for 30 and 60 min, respectively) of wild-type (WT) cells led to significant changes in autophagy markers, with 3.7-and 6.1-fold decreases in SQSTM1/p62 expression, and 2.1-and 1.75-fold increases in LC3II expression, respectively; however, minimal or no significant changes were observed in *PAR1* knockdown cells (Fig. 2Di and Dii). Flow cytometry analysis indicated that *PAR1* knockdown reduced lysosomal activity in HAP-treated MCF7 cells (Fig. 2Ei), whereas WT cells exhibited approximately 10-fold increase in lysosomal activity when treated with 0.5 and 0.75 μg/mL

HAP for 30 min, and 9.4-and 8.4-fold increases when treated with 0.25 and 0.5 μg/mL HAP, respectively, for 60 min (Fig. 2Eii).

### *In silico* analyses of the interaction between PAR1 and the peptides

We have previously reported the interaction between PAR-1 and its cleaved N-terminal sequence, PFISED (generated by HAP-mediated cleavage of PAR-1) a tethered ligand in murine breast cancer cells, as the underlying mechanism of HAP-induced apoptosis [28]. However, the interaction between the human homolog of PAR-1 and its N-terminal amino acid sequence at the similar position (PAFISED) remains uninvestigated.

The physicochemical properties of the peptides, such as drug likeness, pharmacokinetics, and toxicity, were analyzed before docking (Supplementary Table 1). Three-dimensional structures of the peptides were predicted, and the top pose was used as a ligand in Autodock Vina (Supplementary Fig. 2A). Interactions with the native ligand vorapaxar suggested that Val257, Tyr337, Leu258, and Ala349 were important residues for the protein–ligand complex (Fig. 3A). The parameters after re-docking vorapaxar were used to perform docking in Autodock Vina, and CABS-dock was used for blind docking (Fig. 3B). The binding affinities of PAFISED and PFISED with PAR1 were examined. Re-docked Vorapaxar showed a binding affinity of-13.136 kcal/mol with hydrogen atoms of Tyr337 and Val257 of PAR1, while PAFISED and PFISED showed-6.721 and-5.160 kcal/mol binding affinities, respectively (Fig. 3B). PAFISED interacted with hydrogen atoms of Tyr337, Leu263, and Thr261, whereas PFISED interacted with those of Tyr337, Tyr95, Tyr350, and Tyr187 in top most docking pose (Fig. 3B, Ci, and Cii). Both peptides showed considerable interaction with PAR-1 in blind docking. The cluster densities of PAFISED and PFISED were 16.34 and 29.33; average root mean square deviations (RMSDs) were 7.40 and 2.86; and maximum RMSDs were 32.88 and 28.90 with the number of elements in selected cluster being 121 and 84, respectively (Supplementary Fig. 2C and D). The hydrogen bond interactions (Supplementary Table 2) and hydrophobicity (Supplementary Fig. 2B) of the top docked poses by CABS dock were also analyzed by Biovia Discovery Studio visualizer.

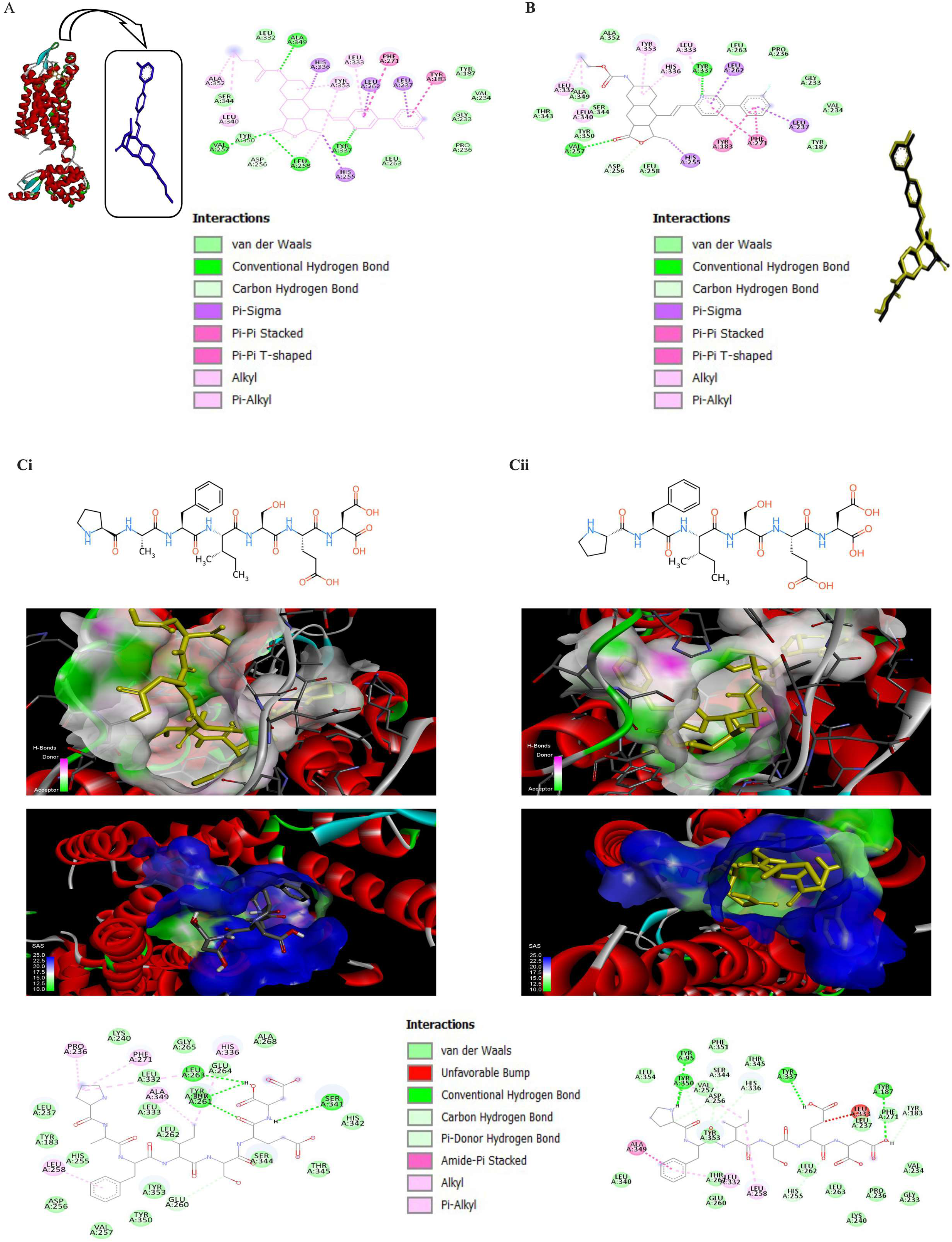

### Noncanonical activation of PAR1 by HAP regulates the induction of autophagy

To compare between thrombin-mediated canonical PAR1 activation and noncanonical activation of PAR1 by HAP regarding their potential of inducing autophagy, we analyzed LC3II and LAMP1 expression in MCF7 and 4T1 cells. No significant upregulation of these markers was observed in thrombin-treated cells compared to that in the control. Conversely, HAP-treated MCF7 cells showed 16.8-and 6.6-fold increases in LC3II and LAMP1 expression, respectively, while 4T1 cells exhibited 8.65-and 3-fold increases in LC3II and LAMP1 expression, respectively. HAP treatment downregulated phosphorylated mTOR (p-mTOR) expression by 4.2-and 6.8-fold in MCF7 and 4T1 cells, respectively, consistent with the notion that mTOR activation negatively regulates autophagy. However, the increase in mTOR phosphorylation in either of the thrombin-treated cells was insignificant (Fig. 4Ai and Aii).

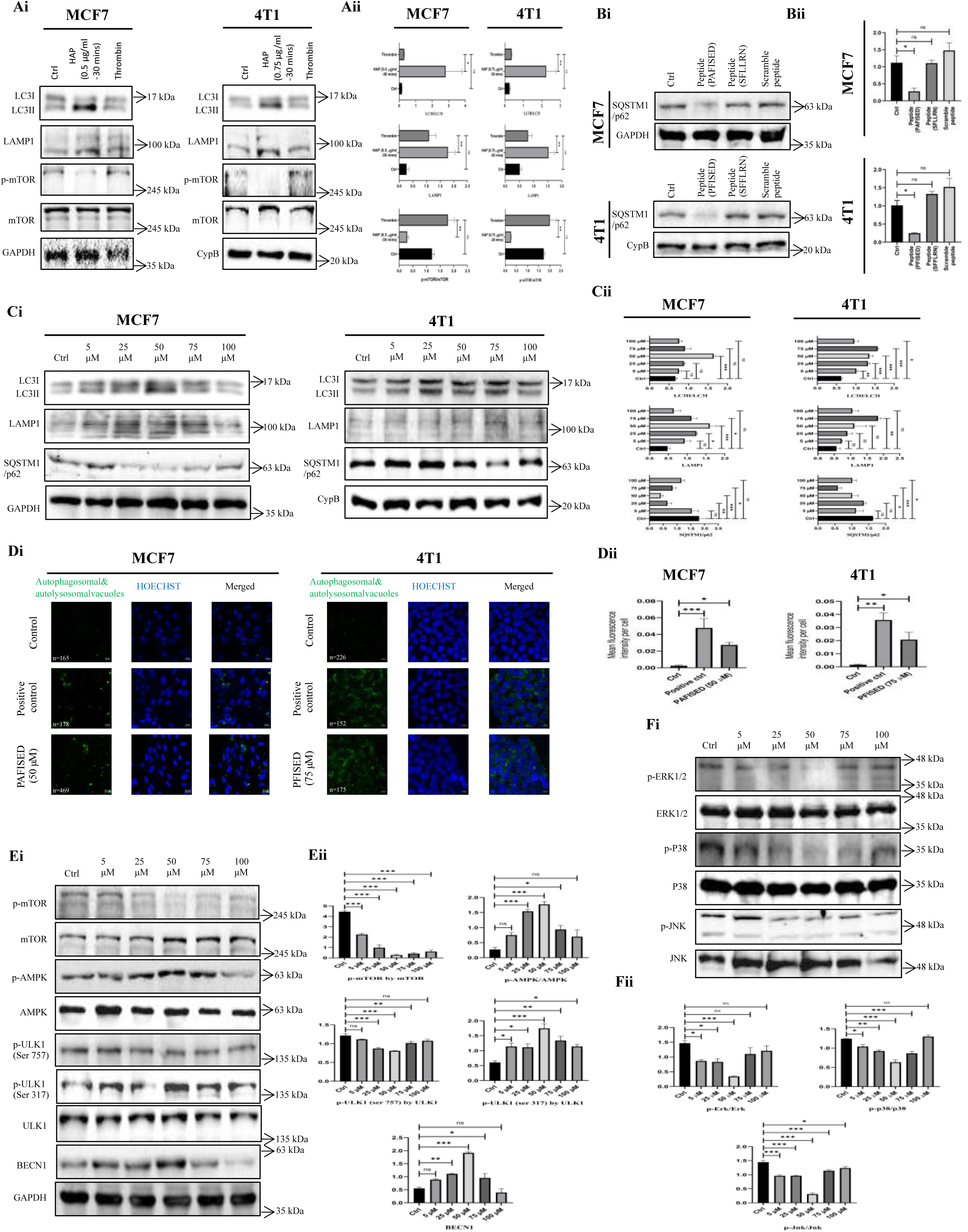

### Potential of the synthetically designed peptides in inducing autophagy

In EAC cells, HAP-mediated cleavage of PAR1 exposes the PFISED sequence, which further binds to PAR1 as a tethered ligand [28]. The human PAR1 homolog (UniProt ID: P25116) possesses a similar sequence, PAFISED, with an additional Ala residue between Pro and Phe at the N-terminal end. *in silico*analysis has shown that PAFISED can bind to PAR1. The synthetic hexapeptide SFLLRN mimics the effect of thrombin on PAR1 without requiring proteolytic cleavage [60]. To explore the autophagic potential of PAFISED and PFISED, we assessed their effects on LC3, LAMP1, and SQSTM1/p62 expression. Optimal autophagy induction was observed with 50 μM PAFISED in MCF7 cells and 75 μM PFISED in 4T1 cells (Fig. 4Ci). Densitometric analysis revealed a 2.47-fold increase in the LC3II/LC3I ratio and a 3.35-fold increase in LAMP1 expression in MCF7 cells. In 4T1 cells, the LC3II/LC3I ratio and LAMP1 expression increased by 2.52-and 3.23-fold, respectively. SQSTM1/p62 expression decreased by 4.4-and 2.76-fold in MCF7 and 4T1 cells, respectively (Fig. 4Cii). Comparisons of PAFISED and PFISED with the thrombin-mimicking peptides SFLLRN (human) and SFFLRN (mouse), and a scramble peptide showed a 4-fold reduction in SQSTM1/p62 expression by PAFISED and PFISED compared to that by their respective controls. No significant change was noted in cells treated with SFLLRN, SFFLRN, or scramble peptides (Fig. 4Bi and Bii). Autophagy induction was further confirmed by confocal microscopy (Fig. 4Di), showing significant increases in AP and AL vacuoles in peptide-treated cells accounting for approximately 12-and 13-fold increases in MFI per cell in MCF-7 and 4T1 cells, respectively (Fig. 4Dii). Flow cytometric analyses using the corroborated these results (Supplementary Fig. 1Ai) showing significant increases of MFI in peptide-treated cells (Supplementary Fig. 1Aii).

Additionally, no significant autophagy induction was observed in normal breast epithelial cells (MCF10A), which showed reduced PAR1 expression compared to that in MCF7 cells, and LC3 and LAMP1 expression was almost negligible (Supplementary Fig. 3C).

### Signaling pathway of peptide-induced autophagy

Immunoblotting revealed that treatment of MCF7 cells with 50 µM PAFISED significantly reduced p-mTOR levels by 16.21-fold, while phosphorylated AMPK levels increased by 6.5-fold compared to that of the control. ULK1 phosphorylation showed a 1.5-fold decrease in phosphorylated ULK (p-ULK1)^Ser757^ and a 2.86-fold increase in p-ULK1^Ser317^ (active form), suggesting AMPK-mediated ULK1 activation, outperforming mTOR-mediated phosphorylation of ULK1 at Ser757 [17]. Notably, Beclin 1 (BECN1) expression increased by 3.46-fold, indicating the recruitment of BECN1 to form BECN1–VPS34–VPS15 core complex that is instrumental for the nucleation phase of autophagy (Fig. 4Ei and Eii). Further analysis revealed > 4-fold decreases in ERK1/2 and JNK phosphorylation, and a 2-fold reduction in p38 phosphorylation, highlighting the molecular mechanisms behind PAR1-mediated mTOR downregulation by PAFISED (Fig. 4Fi and Fii).

### Potential of HAP and the peptides in forming AL

To investigate AP–lysosome fusion, we used pMXs GFP-LC3-RFP plasmid as an autophagy flux reporter. Following transfection, endogenous ATG4 proteins cleave the plasmid, producing GFP-LC3 and red fluorescent protein (RFP) in equimolar amounts. Upon AP– lysosome fusion, GFP-LC3 is quenched in the acidic environment, whereas RFP fluorescence remains intact. An increased RFP:GFP ratio indicates autophagy flux [52]. In MCF7 and 4T1 cells treated with HAP, confocal microscopy revealed 3-and 5-fold in the RFP:GFP ratio, respectively, while PAFISED treatment showed an 8-fold increase in MCF7 cells, and PFISED treatment resulted in a 3-fold increase in 4T1 cells (Fig. 5Ai and Aii).

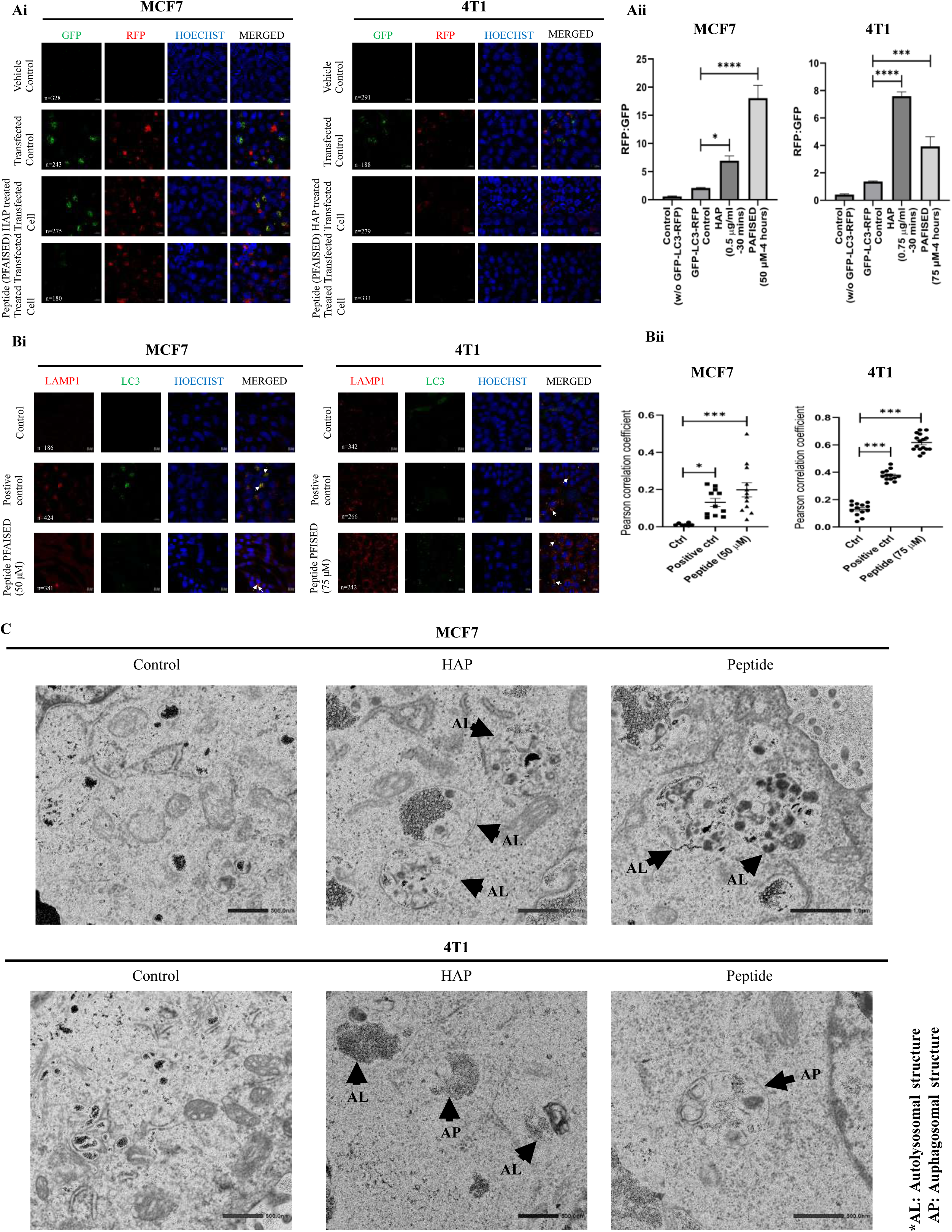

To validate our findings, we assessed the colocalization of LC3 and LAMP1 in peptide-treated MCF7 and 4T1 cells using confocal microscopy (Fig. 5Bi). The Pearson’s correlation coefficient for colocalization increased by 15.5-and 4.7-fold in MCF7 and 4T1 cells, respectively (Fig. 5Bii), indicating the fusion of AP with lysosome. Transmission electron microscopy further confirmed AL formation, characterized by multiple dark patches indicative of degraded cargo, in HAP-and peptide-treated MCF7 and 4T1 cells (Fig. 5C). However, we also observed some AP structures containing intact cellular organelles in HAP-and peptide-treated 4T1 cells.

### Determination of cancer stage of breast tumors developed using 4T1 cells in BALB/c mice

The effect of peptide-induced autophagy on breast cancer cells was validated *in vivo* using a BALB/c mouse model of breast tumor developed by injecting 4T1 cells. Tumor progression, histopathologically assessed weekly over 4 weeks, revealed tumors from weeks 1–3 as Grade I (highly differentiated) and from week 4 as Grade II (moderately differentiated) (Supplementary Fig. 4D and Supplementary Table 3). Tumor grading, based on the Nottingham system [57], correlated with Ki-67 expression. Immunohistochemical analysis revealed a substantial increase in Ki-67 expression in tumors from week 4 compared to that in tumors from weeks 1–3 (Supplementary Fig. 4Ei and Eii), supporting tumor grading based on histopathological analysis. Tumor volume highly increased in weeks 3 and 4 compared to that in weeks 1 and 2 (Supplementary Fig. 4Ci and Cii). Mice showed a slight and nonsignificant decrease in weight until week 3, followed by a slight increase in weight in week 4 (Supplementary Fig. 4B).

### Determination of peptide dose in BALB/c mice

Peptide dose was optimized by assessing peptide-induced toxicity for 2 weeks at an interval of 3 d (Supplementary Fig. 5A). Toxicity was evaluatedby measuring different body parameters of mice, including complete and differential blood counts, alanine aminotransferase (ALT) and aspartate transaminase (AST) activities, blood urea nitrogen (BUN), and creatinine levels (Supplementary Fig. 5B). In a preliminary experiment, the maximum tolerated dose was identified as 20 mg/kg body weight. Beyond this dose, one out of the five mice in a group died, and three started showing visible symptoms of discomfort and itching within 7 d from the time of first administration of peptide (data not shown). In subsequent experiments, mice administered with dosages up to 5 mg/kg body weight showed no notable hepatic toxicity such as increases in ALT and AST activities. Similar results were observed for nephrotoxicity as the increases in creatinine and BUN levels were nonsignificant up to the dosage of 5 mg/kg body weight. However, hemoglobin levels were significantly altered starting from the dosage of 2.5 mg/kg body weight, and the lymphocyte count was altered from the dosage of 1 mg/kg body weight, when compared to those of the control. Eosinophil, monocyte, and neutrophil counts exhibited no significant alteration up to the dosage of 5 mg/kg body weight, consistent with the results of hepatotoxicity and nephrotoxicity (Fig. 7B). Therefore, 5 mg/kg body weight PFISED was optimized for further experiments.

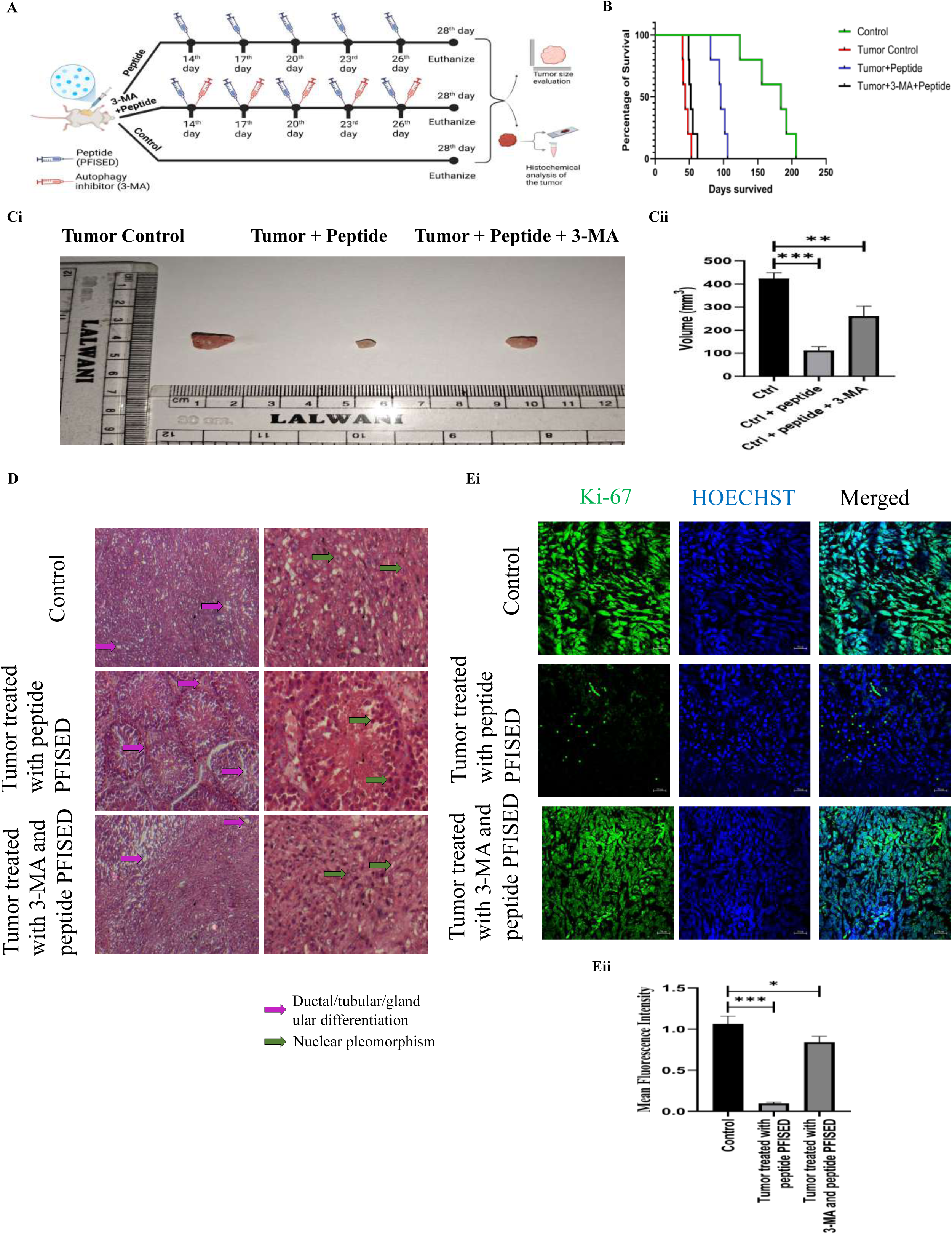

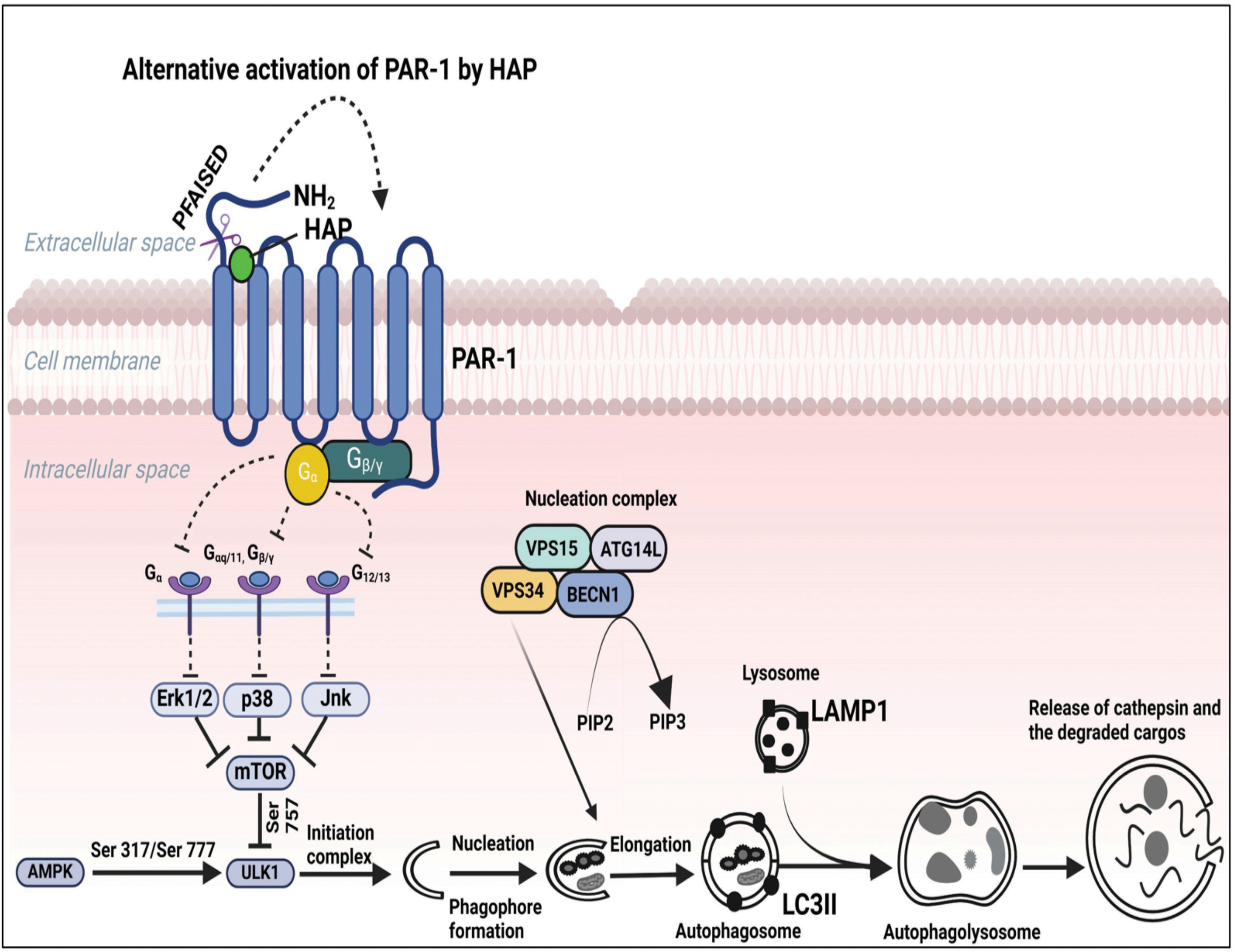

### Peptide-induced autophagy improves survival rate of BALB/c mice carrying low-grade solid breast tumors developed by 4T1 cells

Mice with 2-week-old tumors were used as an early-stage breast cancer model. Untreated mice survived for 45 d on average, whereas PFISED treatment increased the average survival to 96 d. Notably, co-treatment of PFISED and autophagy inhibitor 3-MA reduced the average survival to 54 d. Tumor-free mice lived for approximately 172 d (Fig. 6B).

### Inhibition of tumor growth and progression, and proliferation of tumor cells by peptide-induced autophagy at early stages of breast cancer in BALB/c mice

PFISED treatment resulted in a 3.8-fold reduction of tumor volume in 4T1 cell-injected mice in week 4, compared to that of the control. Conversely, co-treatment of PFISED and 3-MA resulted in only 1.64-fold reduction in tumor volume compared to that of the control (Fig. 6Ci and Cii). Histopathological grading revealed intermediate-grade (II) malignancy for the control and of tumor-bearing mice co-treated with PFISED and 3-MA, whereas in tumor-bearing mice treated with PFISED, low-grade (I) malignancy was noticed (Fig. 6D and Table 1). Ki-67 expression decreased by 10-fold in the PFISED-treated group, whereas it showed only 1.26-fold reduction by co-treatment of PFISED and 3-MA, compared to that of the control group (Fig. 6Ei and Eii).

**Table 1:**
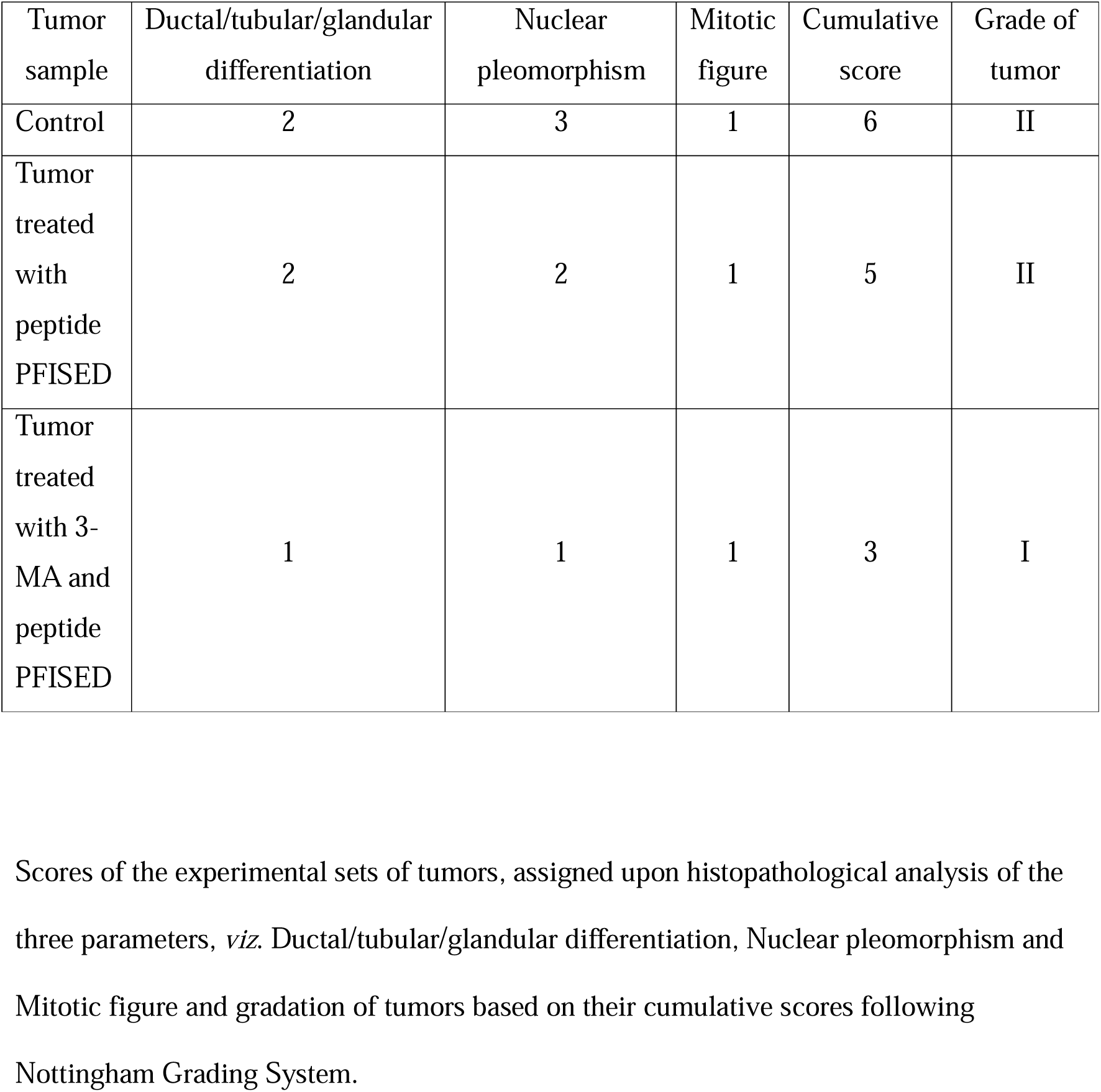
Analysis of the grade of tumors of different experimental groups.

|  |  |  |  |  |  |
| --- | --- | --- | --- | --- | --- |
| Control | 2 | 3 | 1 | 6 | II |
| Tumor treated with peptide PFISED | 2 | 2 | 1 | 5 | II |
| Tumor treated with 3-MA and peptide PFISED | 1 | 1 | 1 | 3 | I |

| Tumor sample | Ductal/tubular/glandular differentiation | Nuclear pleomorphism | Mitotic figure | Cumulative score | Grade of tumor |
| --- | --- | --- | --- | --- | --- |
| Control | 2 | 3 | 1 | 6 | II |
| Tumor treated with peptide PFISED | 2 | 2 | 1 | 5 | II |
| Tumor treated with 3-MA and peptide PFISED | 1 | 1 | 1 | 3 | I |

### Autophagy-driven cell death by HAP and the peptides

Ki-67 expression indicated increased proliferation of MCF7 and 4T1 cells after HAP and peptide treatments. Pretreatment with 3-MA inhibited HAP-and peptide-induced Ki-67 expression (Supplementary Fig. 3Ai and Aii).

In MCF7 cells, HAP and PAFISED treatments reduced survival by approximately 60% and 50%, respectively (Supplementary Fig. 3Bi), while the survival of 4T1 cells decreased by approximately 46% and 55%, respectively, by HAP and PFISED treatment (Supplementary Fig. 3Bii). Autophagy inhibition by 3-MA pretreatment prior to HAP and peptide treatments increased cell survival.

## Discussion

Autophagy, long regarded as a fundamental cellular self-survival mechanism, has recently garnered relatively high attention for its role in facilitating cell death as well, termed as type II programmed cell death [61–63]. This newfound understanding has led to the exploration of autophagy as a therapeutic target for various diseases, including cancer and neurodegenerative conditions, in which selective elimination of specific cell populations is desired. Autophagy can be induced by different stimuli, indicating the presence of multiple upstream signaling pathways that may govern autophagy independently or in an intertwined manner [2]. In this study, we investigated the potential of a secreted metalloprotease, HAP, in inducing autophagy selectively in mammalian breast cancer cells, alongside the underlying signaling mechanisms.

HAP, owing to its proteolytic activity, engages with PARs on the membranes of various mammalian cancer cells. HAP mediates the alternative activation of PAR1, a key mechanism driving HAP-induced apoptosis. Among the four known subtypes of PARs, HAP specifically binds to PAR1 and activates it [27]. In the present study, HAP induced autophagy in mammalian breast cancer cells at lower concentrations than those previously optimized for apoptosis induction [26]. A single stimulus can simultaneously regulate apoptotic and autophagic pathways, with these processes often acting as mutually exclusive events [29, 30]. We identified that noncanonical activation of PAR1 by HAP was pivotal for the induction of autophagy. When apoptosis and autophagy are triggered by a common stimulus, autophagy is typically initiated as an early response, with apoptosis being recruited at later time points. In some instances, autophagy may even lead to apoptosis [29, 64].

PAR1 is a transmembrane G protein-coupled receptor, which, upon activation, interacts with various G protein subtypes, including G12/13, Gi, and Gq, depending on the cellular context and modulates different signaling pathways [65–69]. Although PAR1 expression is minimal in most noncancerous mammalian cells (with the exception of platelets), it is abundantly expressed in breast cancer cells, making it an ideal target for therapeutic intervention [70]. Thrombin, a serine protease, is a known activator of PAR1. It interacts with the transmembrane receptor, PAR1 through its N-terminal exodomain, which contains two thrombin-binding sites [71]. Thrombin activates PAR1 by cleaving the Arg 41/Ser 42 site of its exodomain [60, 72]. G protein-coupled receptors activate MAP kinases by signaling through G proteins and facilitate proliferation by mTOR activation [73]. This canonical pathway of PAR1 activation by thrombin also escalates proliferation via MAPK activation and leads to mTOR activation [22, 74], which is considered as a negative regulator of autophagy [75]. Our findings suggested that mTOR activation was consistent with thrombin-induced canonical activation of PAR1. However, we observed that HAP treatment downregulated mTOR activation, in contrast to the effects of thrombin. Moreover, thrombin did not induce significant autophagic responses, whereas HAP demonstrated a marked ability of inducing autophagy. To the best of our knowledge, this is the first report that noncanonical PAR1 activation by HAP induces autophagy, as opposed to its canonical thrombin-mediated activation, which primarily upregulates mTOR activation that leads to cell proliferation.

Thrombin activates the human PAR1 homolog by proteolytically cleaving its N-terminal extracellular domain, which exposes a tethered ligand sequence (SFLLRN) that intramolecularly binds PAR1, initiating downstream signaling in platelets and vascular endothelial cells [72]. The murine homolog of PAR1 contains a similar sequence (SFFLRN) with the alteration of one amino acid (Phe44 instead of Leu44) at the same position. HAP cleaves the murine homolog of PAR1 at the Pro90/Pro91 site, thereby generating a new N-terminal end with the sequence PFISED [27]. This sequence is analogous to PAFISED in the human PAR1 homolog, located between Leu84 and Ala92. Both PFISED and PAFISED, designed on the basis of last seven and six amino acids of the N-terminal cleaved ends of human and murine homologs of PAR1, respectively, possess high affinity for their respective PAR1 homologs, as shown by molecular docking studies. The docking result and free energy values suggest that these peptides are stable and potential candidates for cancer treatment.

These peptides efficiently induced autophagy in breast cancer cells in a dose-dependent manner, whereas SFLLRN and SFFLRN failed to induce autophagy. Further investigation into the signaling pathways involved in autophagy induction by PAFISED in MCF7 cells revealed a significant downregulation of mTOR activation, as evidenced by a decrease in p-mTOR levels, which resulted in a substantial decrease in p-ULK1^Ser757^ expression.

Collectively, the major three MAP kinases ERK1/2, p38, and JNK play a notable role in modulating mTOR activation. Given that mTOR is recognized as the master regulator of autophagy, we investigated the effects of PAFISED on the activation of these three MAP kinases. Significant downregulation of the phosphorylation of all three MAP kinases justified reduced p-mTOR expression. However, PAFISED upregulated AMPK phosphorylation, which increased p-ULK1^Ser317^ expression, thereby facilitating the formation of initiation complex. An increased BECN1 expression upon peptide treatment suggested the formation of nucleation complex. As the autophagy-inducing potential of the synthetic peptide that resembled the cleaved N-terminal end of noncanonically activated PAR1 was established, clearly, the induction of autophagy was not HAP-specific, rather it depended on the site of cleavage on the PAR1 exodomain.

A major limitation of many autophagy-inducing agents, including chloroquine, is their inability to form ALs or effectively degrade cargo, an event necessary for autophagy-mediated cell death. Our study confirmed that treatments with both HAP and peptides resulted in the formation of ALs in breast cancer cells. Degradation of cargoes is marked by the rupture of autolysosomal structure, which was evidenced by the elevated expression of Cathepsin L in cytosol. A downregulation of SQSTM1/p62 expression also confirmed AP degradation of cargoes.

In the context of cancer therapy, ensuring that autophagy induction leads to cessation of cellular proliferation and a subsequent reduction in cancer cell survival is crucial. Our data showed a significant decline in both the proliferation and survival of breast cancer cells upon autophagy induction by HAP and the peptides. Notably, breast cancer cells, which exhibit high expression of PAR1, were selectively targeted by HAP and the peptide for autophagy induction, while noncancerous human breast epithelial cells were unaffected.

We also assessed the *in vivo* impact of peptide-induced autophagy in low-grade breast cancer. Using the Nottingham Grading System, we classified tumors based on the assigned scores considering three aspects of the tumor tissue, including ductal/tubular/glandular differentiation, nuclear pleomorphism, and mitotic figure, and correlated them with Ki-67 expression. A cumulative score between 3–5 refers to a low-grade/grade I (well-differentiated) breast cancer, whereas a cumulative score between 6–7 indicates an intermediate-grade/grade II (moderately differentiated) breast cancer. A cumulative score between 8–9 signifies a high-grade/grade III (poorly differentiated) breast cancer [57]. Our findings revealed that breast tumors developed in BALB/c mice by 4T1 cells remained well-differentiated (Grade I) up to 3 weeks. Following PFISED treatment, we observed significant inhibition of tumor growth and progression along with a partial cessation of increasing proliferation rate in this early-stage, low-grade breast cancer model. Additionally, the peptide was safe to be administered up to the dosage of 5 mg/kg body weight. However, further studies on the physicochemical properties of the peptide and large-scale *in vivo* experiments in higher order animals including primates are warranted before advancing to extensive and further clinical trials.

Although our study demonstrated that noncanonical activation of PAR1 inhibits tumor growth, a significant limitation lies in the lack of investigation into its effects on platelet activation. Considering the role of canonical PAR1 activation by thrombin in platelet activation, assessment of the impact of this noncanonical activation of PAR1 on platelets is imperative to rule out any potential thrombotic complication prior to any consideration of clinical translation. As this aspect was beyond the purview of the current study, we intend to address it in future research.

In conclusion, this study opens new avenues for therapeutic strategies aimed at leveraging autophagy as a treatment modality for breast cancer, specifically in the early stage.

## Glossary

3-MA: 3-Methyladenine
4T1: 410.4 subpopulation Tumor 1
AL: Autolysosome
ALT: Alanine transaminase
AP: Autophagosome
AST: Aspartate aminotransferase
BUN: Blood urea nitrogen
EAC: Ehrlich ascites carcinoma
GABARAP: Gamma-aminobutyric acid receptor-associated protein
HAP: Hemagglutinin protease
KD: Knocked down
MCF10A: Michigan cancer foundation-10A
MCF7: Michigan cancer foundation-7
MEM: Minimum essential medium
PAR1: Protease activated receptor 1
PARs: Protease activated receptors
RPMI 1640: Roswell park memorial institute 1640
TEM: Transmission electron microscope
WT: Wild type

## Declaration of generative AI

During the preparation of this work the authors used ChatGPT-4 in order to modify and improve the quality of language in some parts of the manuscript. After using this tool, the authors reviewed and edited the content as needed and take full responsibility for the content of the publication.

## Data Statement

The data presented in this study are available in the article and in the supplementary information. Any additional information required to reanalyse the data reported in this paper is available from the lead contact upon request. Information and requests for resources should be directed to and will be fulfilled by the lead contact, Dr. Amit Pal.

## Author contributions

S.S. conceptualized and designed the study, performed experiments, analyzed and validated the data, wrote the original draft and reviewed. R.K. performed the *in silico* experiments and analyzed the data. S.S. analyzed the histopathological data. A.G. provided technical assistance in confocal microscopy experiments. A.P. helped in flow cytometric data acquisition and formal analysis. R.T. did formal analysis and assisted in microbiological experiments. N.N. did formal analysis. A.P. conceptualized the study, supervised the research work, helped in funding acquisition, reviewed and approved the final manuscript.

## Acknowledgements

Mr. Saibal Saha extends his sincere gratitude to the Indian Council of Medical Research for providing financial support for this research and to the University Grants Commission, India, for awarding him the esteemed UGC-NET-JRF fellowship (UGC Ref. No.: 529/CSIR-UGC-NET JUNE 2019).

We are truly grateful to Dr. Santasabuj Das, Director of ICMR-NIRBI, for granting access to the central research instrumentation facility at ICMR-NIRBI. Our heartfelt thank goes to Dr. Sushmita Bhattacharya (ICMR-NIRBI, India) for her invaluable inputs and suggestions in this study. We are also thankful to Dr. Somsubhra Nath (Presidency University, Kolkata, India) and Dr. Saptak Banerjee (CNCI, Kolkata, India) for their generous provision of cell lines.

We would like to take the opportunity to express our gratitude to Dr. Sneha Mitra (ICMR-NIRBI, India) for her invaluable assistance in the analysis of flow cytometry data, and to Dr. Sarthak Das (A*STAR, Singapore) for his technical help in figure formatting. We also wish to acknowledge Mrs. Arpita Sarbajna (ICMR-NIRBI, India) and Ms. Anaswara Ramesh (ICMR-NIRBI, India) for their essential support during the transmission electron microscopy experiments.

