## Supplementary data for "Noncanonical activation of protease-activated receptor 1 induces autophagy in mammalian breast cancer cells"

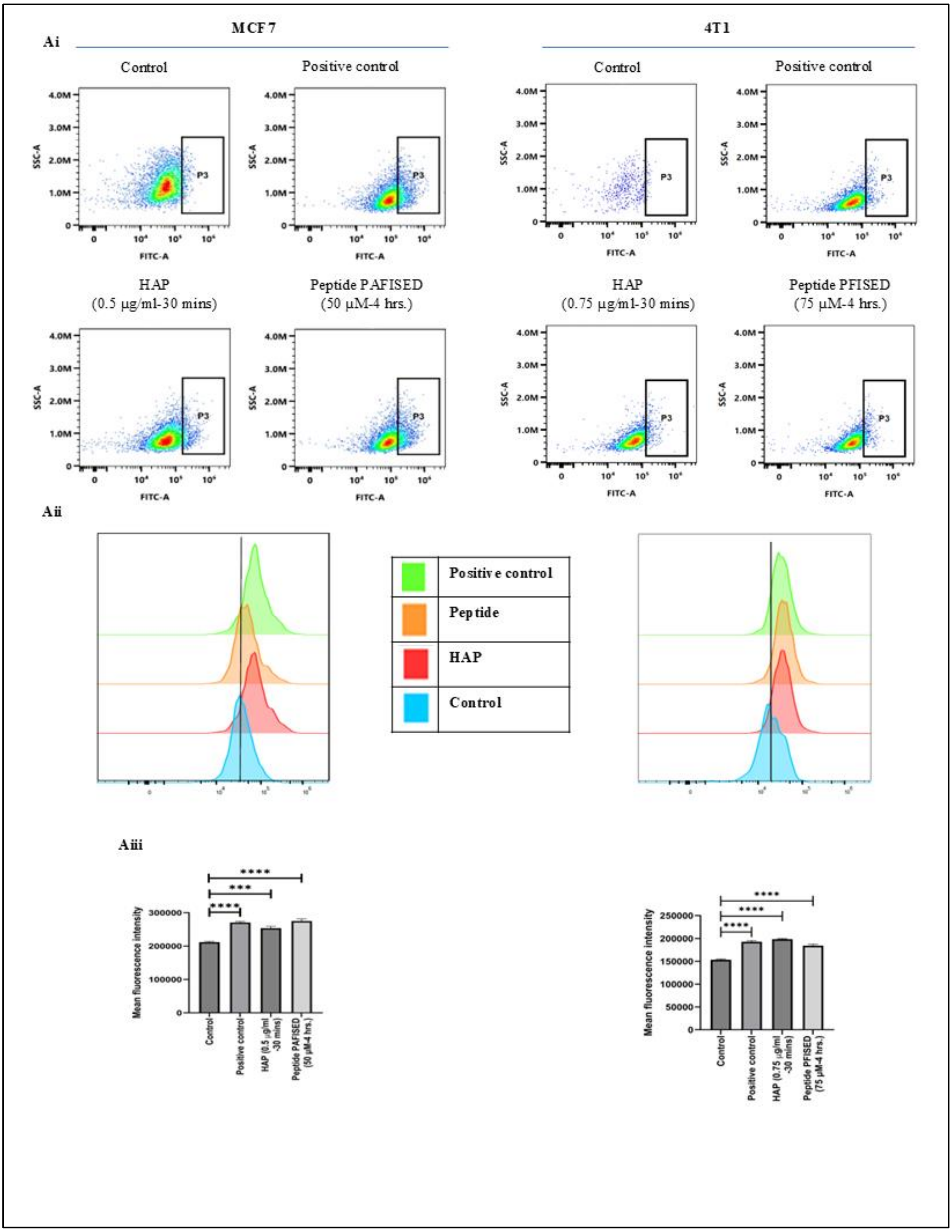
Ai
MCF7
4T1
Control
Positive control
HAP
(0.5 µg/ml-30 mins)
Peptide PAFISED
(50 µM-4 hrs.)
HAP
(0.75 µg/ml-30 mins)
Peptide PFISED
(75 µM-4 hrs.)
SSC-A
FITC-A
P3
Aii
Positive control
Peptide
HAP
Control
Aiii
Mean fluorescence intensity
Control
Positive control
HAP (0.5 µg/ml -30 mins)
Peptide PAFISED (50 µM-4 hrs.)
HAP (0.75 µg/ml -30 mins)
Peptide PFISED (75 µM-4 hrs.)

**Figure S1: Autophagy assay analysis. (Ai)** Representative flowcytometric images of MCF7 and 4T1 cells treated with HAP (0.5 and 0.75 μg/ml for MCF-7 and 4T1 respectively) for 30 minutes and with the peptides (50 μM of PAFISED for MCF-7 and 75 μM of PFISED for 4T1) for 4 hours and stained using the Autophagy assay kit (ab139484) of Abcam. Cells treated with rapamycin and chloroquine simultaneously for 24 hours was kept as positive control. Fluorescence was measured under FITC channel of a flowcytometer. All the representative histograms in each cell type are arranged vertically for a better understanding of comparative analysis. **(Aii)** Graphical analysis of Mean fluorescence intensity of FITC.

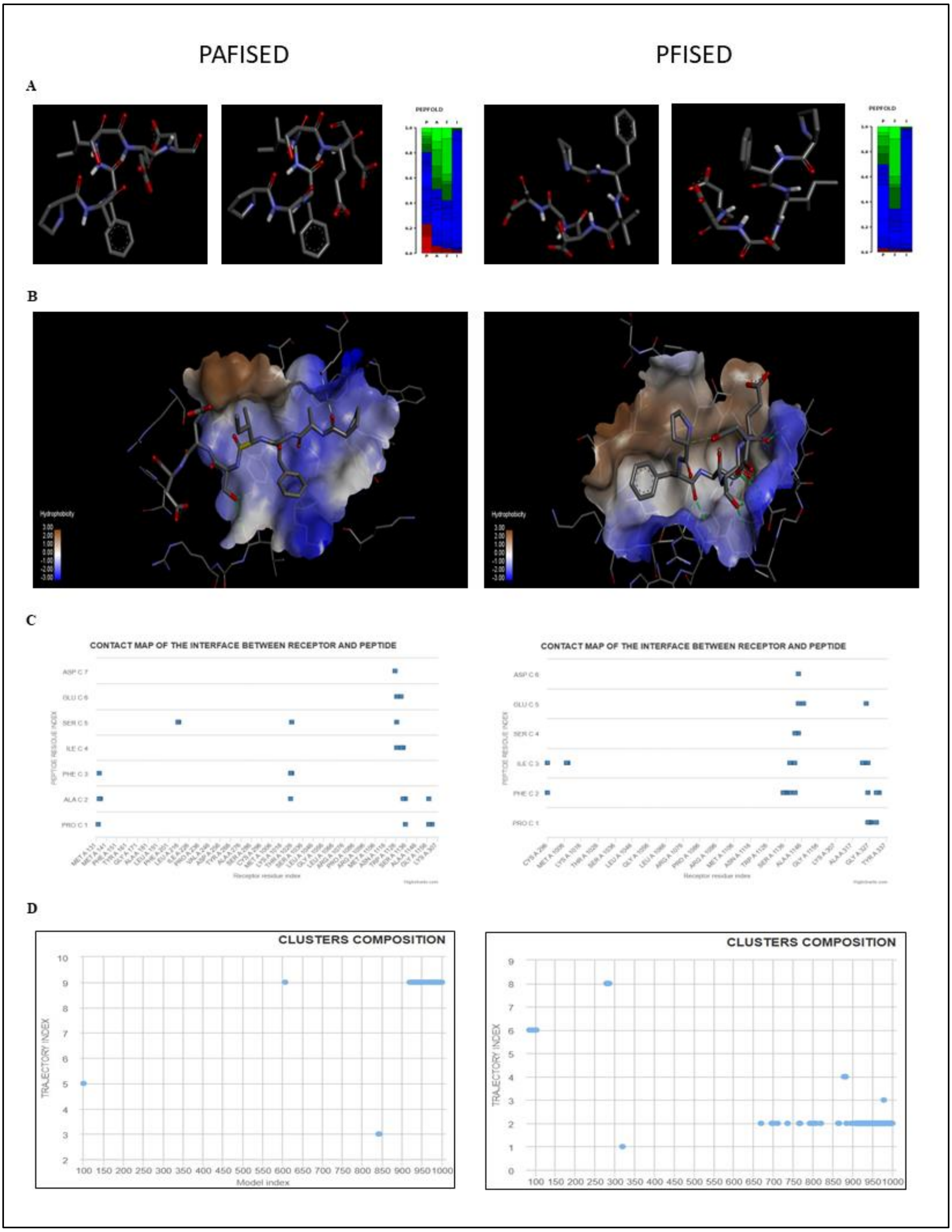
PAFISED
PFISED
A
B
C
D
CONTACT MAP OF THE INTERFACE BETWEEN RECEPTOR AND PEPTIDE
CONTACT MAP OF THE INTERFACE BETWEEN RECEPTOR AND PEPTIDE
CLUSTERS COMPOSITION
CLUSTERS COMPOSITION
TRAJECTORY INDEX
Model index

**Figure S2: Peptide Structure prediction and Blind Docking Analysis of the peptides PAFISED (Left Side) & PFISED (Right side). (A)** Top 2 predicted peptide structure by PEP-FOLD4 with local structure prediction profile which shows the probabilities of structural alphabet. **(B)** hydrophobicity representation of peptide with PAR1 by CABS dock. **(C)** Contact Map of receptor residues and Peptide of top docked pose. **(D)** Cluster composition vs trajectory affiliations of top docked pose.

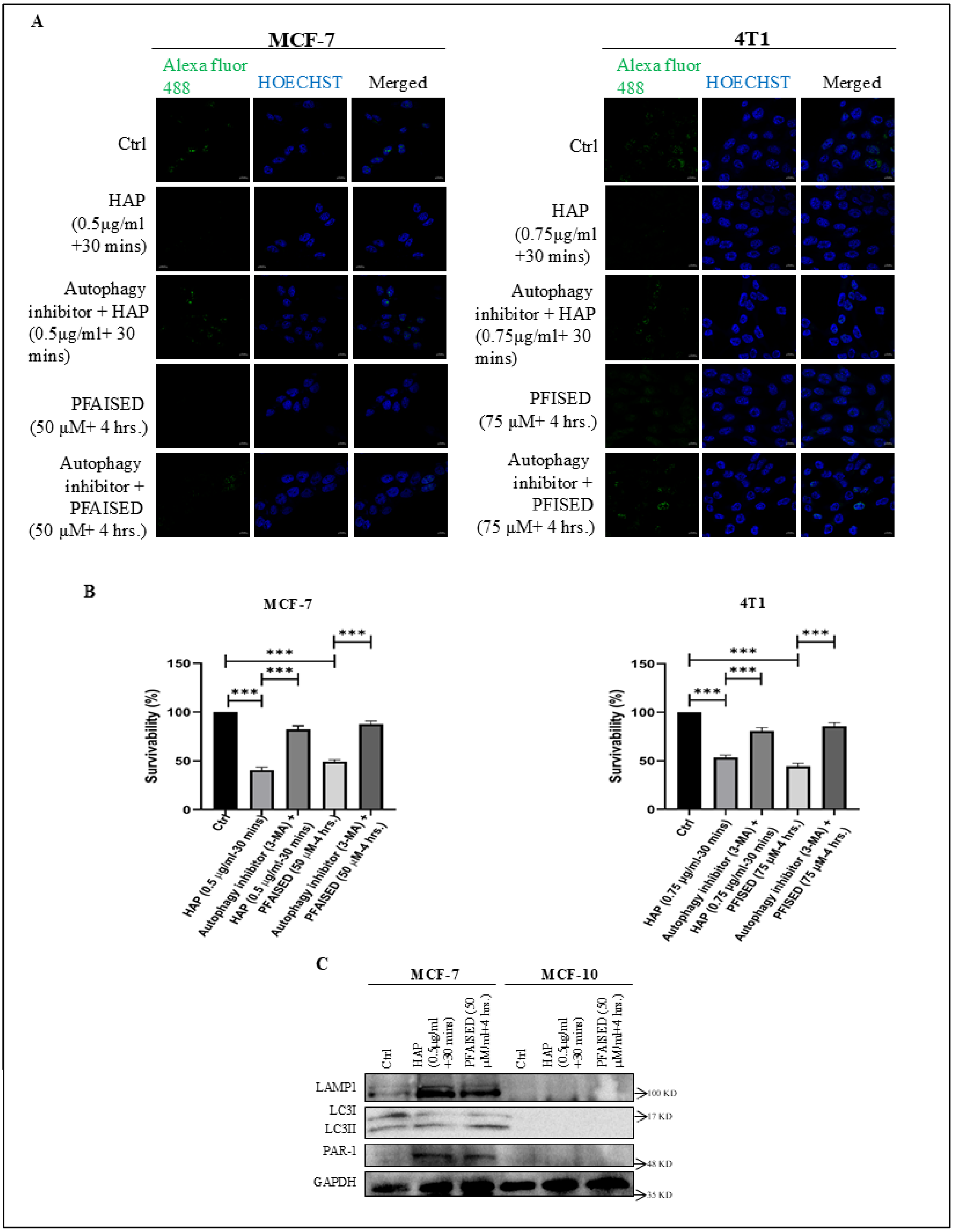
A
MCF-7
Alexa fluor 488
HOECHST
Merged
Ctrl
HAP (0.5µg/ml +30 mins)
Autophagy inhibitor + HAP (0.5µg/ml+ 30 mins)
PFAISED (50 µM+ 4 hrs.)
Autophagy inhibitor + PFAISED (50 µM+ 4 hrs.)
4T1
Alexa fluor 488
HOECHST
Merged
Ctrl
HAP (0.75µg/ml +30 mins)
Autophagy inhibitor + HAP (0.75µg/ml+ 30 mins)
PFISED (75 µM+ 4 hrs.)
Autophagy inhibitor + PFISED (75 µM+ 4 hrs.)
B
MCF-7
Survivability (%)
150
100
50
0
***
Ctrl
HAP (0.5 µg/ml-30 mins)
Autophagy inhibitor (3-MA) + HAP (0.5 µg/ml-30 mins)
PFAISED (50 µM-4 hrs.)
Autophagy inhibitor (3-MA) + PFAISED (50 µM-4 hrs.)
4T1
Survivability (%)
150
100
50
0
***
Ctrl
HAP (0.75 µg/ml-30 mins)
Autophagy inhibitor (3-MA) + HAP (0.75 µg/ml-30 mins)
PFISED (75 µM-4 hrs.)
Autophagy inhibitor (3-MA) + PFISED (75 µM-4 hrs.)
C
MCF-7
MCF-10
Ctrl
HAP (0.5µg/ml +30 mins)
PFAISED (50 µM/ml+4 hrs.)
Ctrl
HAP (0.5µg/ml +30 mins)
PFAISED (50 µM/ml+4 hrs.)
LAMP1
LC3I
LC3II
PAR-1
GAPDH
100 KD
17 KD
48 KD
35 KD

**Figure S3: Effect of HAP and peptide induced autophagy on the proliferation and viability of cancer cells: (A)** Representative immunofluorescence images of MCF-7 and 4T1 cells treated with HAP (0.5 and 0.75 μg/ml for MCF-7 and 4T1 respectively) for 30 minutes and with the peptides (50 μM of PAFISED for MCF-7 and 75 μM of PFISED for 4T1) for 4 hours with or without pre-treatment of autophagy inhibitor 3-MA and analyzed for proliferation using anti-Ki-67 antibody and AlexaFluor 488 tagged corresponding secondary antibody. Nuclei were stained with Hoechst 33342. Images were captured using a confocal microscope. **(B)** Survivability analysis of MCF-7 and 4T1 cells treated with HAP (0.5 and 0.75 μg/ml for MCF-7 and 4T1 respectively) for 30 minutes and with the peptides (50 Mm of PAFISED for MCF-7 and 75 μM of PFISED for 4T1) for 4 hours with or without pre-treatment of autophagy inhibitor 3-MA by MTT assay. Absorbance was measured using a spectrophotometric plate reader and survivability was calculated considering the control cells have a 100% survivability. The survivability percentages are presented graphically. **(C)** Immunoblots of LAMP1, LC3 and PAR-1 in the whole cell lysates of MCF7 and MCF10A, treated with HAP (0.5 μg/ml for 30 minutes) and the peptide PAFISED (50 μM for 4 hours).

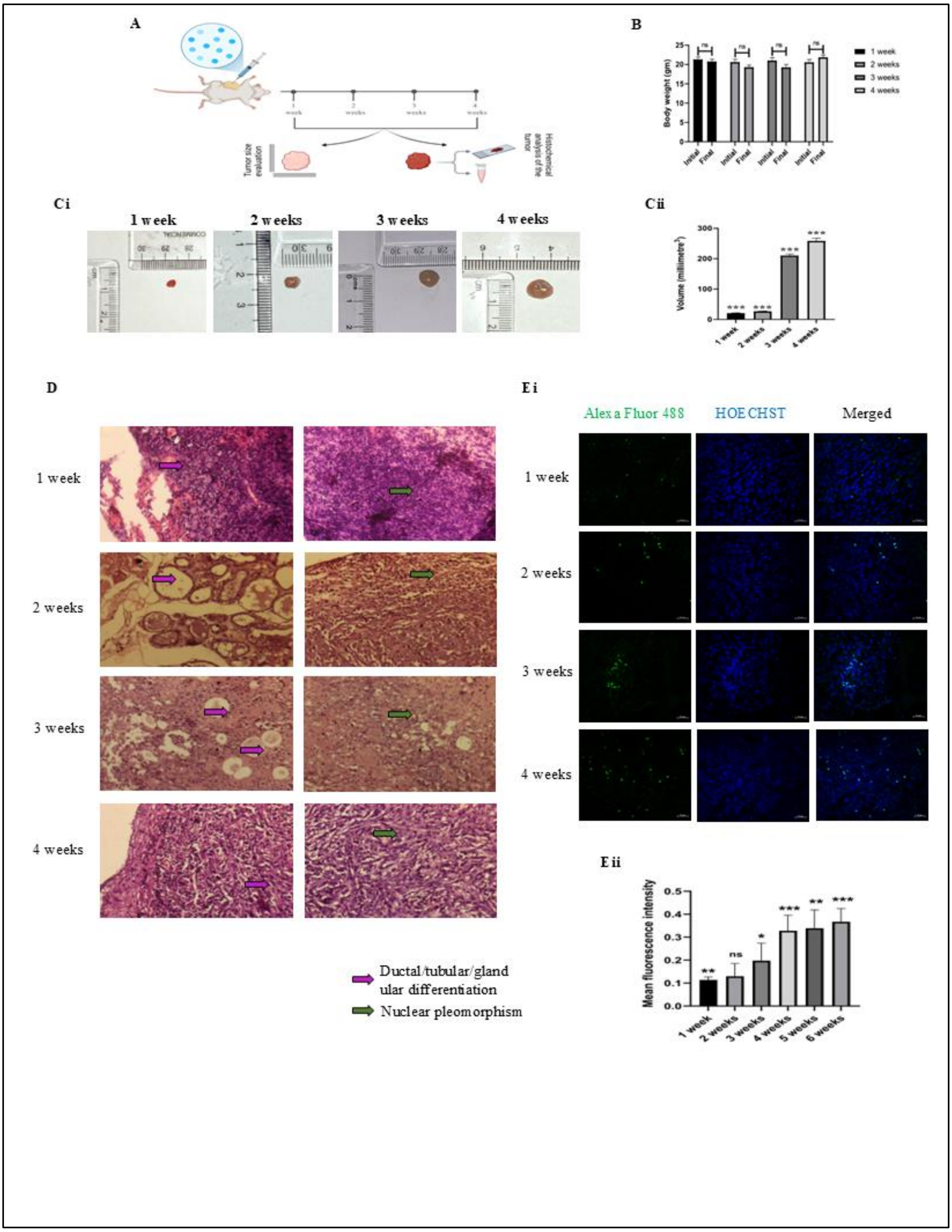
A
Tumor size evaluation
Histochemical analysis of the tumor
B
Body weight (gm)
Initial
Final
ns
1 week
2 weeks
3 weeks
4 weeks
Ci
1 week
2 weeks
3 weeks
4 weeks
Cii
Volume (millimetre³)
D
1 week
2 weeks
3 weeks
4 weeks
Ductal/tubular/glandular differentiation
Nuclear pleomorphism
Ei
Alexa Fluor 488
HOECHST
Merged
1 week
2 weeks
3 weeks
4 weeks
Eii
Mean fluorescence intensity
1 week
2 weeks
3 weeks
4 weeks
5 weeks
6 weeks

**Figure S4: Gradation of murine breast cancer. (A)** Schematic diagram of the outline of the experiments done to determine the grades of breast cancer developed by 4T1 cells in BALB/c mice upto six weeks. **(B)** Graphical presentation of body weights of mice with breast cancer at each week's interval upto 6 weeks starting from the day of injecting 4T1 cells. **(Ci)** Images of breast tumors developed by 4T1 cells in BALB/c mice. **(Cii)** Graphical comparison of the volumes of tumors of different time points. **(D)** Representative histological images of tumors of different time points with H&E staining indicating ductal/ tubular/ glandular differentiation (purple arrow) and nuclear pleomorphism (green arrow). **(Ei)** Representative immunohistochemical images of tumor tissues of different time points stained against Ki-67. **(Eii)** Quantification of mean fluorescence intensity of Ki-67, normalized against nuclear staining with Hoechst 33342.

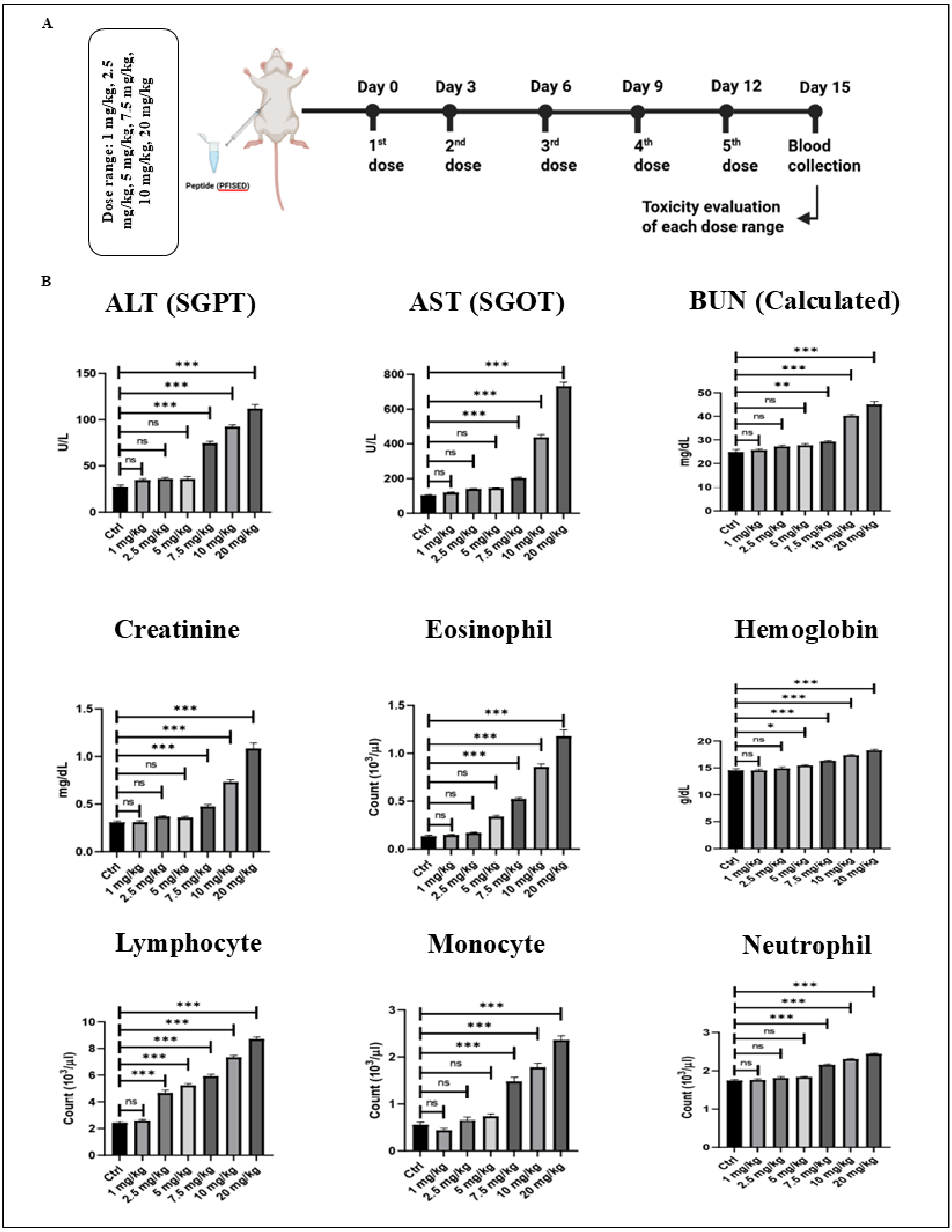
A
Dose range: 1 mg/kg, 2.5 mg/kg, 5 mg/kg, 7.5 mg/kg, 10 mg/kg, 20 mg/kg
Peptide (PFISED)
Day 0
Day 3
Day 6
Day 9
Day 12
Day 15
1st dose
2nd dose
3rd dose
4th dose
5th dose
Blood collection
Toxicity evaluation of each dose range
B
ALT (SGPT)
AST (SGOT)
BUN (Calculated)
Creatinine
Eosinophil
Hemoglobin
Lymphocyte
Monocyte
Neutrophil
U/L
mg/dL
g/dL
Count (10³/µl)
Ctrl
1 mg/kg
2.5 mg/kg
5 mg/kg
7.5 mg/kg
10 mg/kg
20 mg/kg

**Figure S5: (A)** Schematic diagram of the design of the experiment for toxicity evaluation of the peptide PFISED in BALB/c mice. (B) Assessment of hepatotoxicity parameters (ALT and AST), nephrotoxicity parameters (creatinine and calculated BUN) and hematological parameters (eosinophil, hemoglobin, lymphocyte, monocyte and neutrophil) in BALB/c mice given peptide of different dosages ranging from 1 mg/kg body weight to 20 mg/kg body weight at an interval of 3 days for 12 days.

**Table S1: The physico-chemical properties of Peptide checked by SwissADME**

| ADMET | PAFISED | PFISED |
|---|---|---|
| Formula | C35H51N7O13 | C32H46N6O12 |
| Molecular Weight | 777.82 g/mol | 706.74 g/mol |
| H-bond Acceptors | 14 | 13 |
| H-bond Donors | 11 | 10 |
| Lipinski | No | No |
| Veber | No | No |
| Bioavailability | 0.11 | 0.11 |
| GI Absorption | Low | Low |
| Water Solubility | Soluble | Soluble |
| Synthetic Accessibility | 6.46 | 5.95 |
| Blood-Brain Barrier Permeant | No | No |
| Toxicity | No | No |

An analysis of the drug-like properties, pharmacokinetics, toxicity, and physicochemical properties of the peptides

**Table S2: The hydrogen bond interactions of top docked pose by CABS-dock**

| | PAFISED | PFISED |
|---|---|---|
| Hydrogen Bond | A:LYS135:NZ - C:ALA2:O | A:THR1142:OG1 - C:PHE2:O |
| | A:TYR1025:OH - C:SER5:OG | A:ASN1144:ND2 - C:SER4:OG |
| | C:SER5:OG - A:LEU216:O | A:ASN1144:ND2 - C:SER4:O |
| | C:GLU6:N - A:ASP1127:OD2 | A:ASN1144:ND2 - C:ASP6:O |
| | A:LYS1135:CE - C:PRO1:O | A:LYS1147:NZ - C:GLU5:O |
| | | C:ILE3:N - A:CYS296:SG |
| | | C:SER4:OG - A:THR1142:O |
| | | A:ARG1137:CD - C:PHE2:O |
| | | C:PRO1:CA - C:ASP6:OD1 |
| | | C:SER4:CB - C:ASP6:O |

The hydrogen bond interactions between receptor residue and peptides of top docked pose by CABS-dock. The interactions were analysed by Biovia Discovery Studio visualizer.

**Table S3: Identification of low-grade breast carcinoma by gradation of tumor**

| Tumor sample | Ductal/tubular/glandular differentiation | Nuclear pleomorphism | Mitotic figure | Cumulative score | Grade of tumor |
|---|---|---|---|---|---|
| 1 week | 2 | 1 | 1 | 4 | I |
| 2 weeks | 1 | 2 | 1 | 4 | I |
| 3 weeks | 2 | 2 | 1 | 5 | I |
| 4 weeks | 3 | 2 | 1 | 6 | II |

Scoring pattern of the tumors harvested at four different time points. Scores were attributed upon histopathological analysis of the three parameters, *viz*. Ductal/tubular/glandular differentiation, Nuclear pleomorphism and Mitotic figure and gradation of tumors based on their cumulative scores according to Nottingham Grading System.
